# ATM promotes reversed fork processing during DNA interstrand cross-link repair

**DOI:** 10.1101/2025.10.08.680157

**Authors:** Maria Altshuller, Victoria A. MacKrell, Jasmine Tzeng, Richa Nigam, Ting-Yu Wang, Tsui-Fen Chou, Daniel R. Semlow

## Abstract

During replication-coupled DNA interstrand cross-link (ICL) repair, fork reversal is thought to enable the Fanconi anemia (FA) pathway to resolve the ICL through nucleolytic incisions. Subsequent fork restoration then allows nascent DNA strand extension past the lesion. Although these fork remodeling events are crucial for ICL repair, their regulation remains poorly understood. Here, we use cell-free *Xenopus* egg extracts to investigate fork dynamics during ICL repair by the FA pathway. We find that the ataxia telangiectasia-mutated (ATM) kinase is activated concomitantly with fork reversal and promotes resection of the reversed fork intermediate. This resection depends on the coordinated activities of the EXO1 and DNA2 nucleases. Our data indicate that EXO1 initiates 5’ to 3’ resection of nascent lagging strands in the regressed arm, while DNA2 performs 5’ to 3’ resection of recessed lagging strands. We further show that the inhibition of protein phosphatase 2A (PP2A) during ICL repair results in ATM hyperactivation, reversed fork over-resection, and formation of aberrant end-joining products, indicating that PP2A counteracts ATM signaling to constrain reversed fork resection. Taken together, this work implicates reversed forks as substrates for ATM activation and reveals a phospho-regulatory circuit that governs reversed fork processing during ICL repair.

## Introduction

DNA interstrand cross-links (ICLs) covalently link the two strands of DNA and thereby block strand separation. These lesions are produced by endogenous cellular metabolites (e.g. formaldehyde, crotonaldehyde, abasic sites), microbiome-derived secondary metabolites (e.g. colibactin), and chemotherapeutics (e.g. cisplatin, nitrogen mustard derivatives, psoralens)^1,2^. Mutations in genes associated with ICL repair cause human diseases including the chronic kidney disease karyomegalic intestinal nephritis and the bone marrow failure and cancer predisposition syndrome Fanconi anemia. ICLs represent formidable impediments to DNA replication and can lead to deletions, chromosomal rearrangements, and chromosome missegregation. Consequently, even a single unrepaired ICL can be lethal to a eukaryotic cell^3^.

Among the multiple mechanisms that cells have evolved to resolve ICLs, the Fanconi anemia (FA) pathway is thought to be the most important for ICL resistance in proliferating cells^1^. The FA pathway is coupled to DNA replication and can be triggered by the convergence and stalling of replisomes on either side of an ICL (Fig. 1a)^4^. Fork convergence induces TRAIP-dependent ubiquitylation and p97-dependent unloading of the replicative CDC45-MCM2-7-GINS (CMG) helicase^5–7^. One of the two forks abutting the ICL undergoes extensive reversal, in which reannealing of the parental DNA strands leads to the displacement and annealing of nascent strands and the formation of a four-helix junction^8^. Interestingly, during ICL repair in *Xenopus* egg extracts, this fork reversal is coupled to resection of the nascent lagging strand resulting in a regressed arm that comprises predominantly single-stranded DNA^8^. Although a requirement for fork reversal during ICL repair has not been proven, reversed forks appear to represent on pathway ICL repair intermediates. Additionally, fork reversal establishes the ICL within a duplex at a three-helix junction. This configuration represents the preferred substrate for the XPF-ERCC1 3’ flap endonuclease, which incises one parental DNA strand 5’ of the ICL^9^. A second nuclease, such as SΝΜ1A, mediates 5’ to 3’ expansion of the XPF-induced nick past the ICL to release (“unhook”) the cross-link and generate a DNA double strand break (DSB). Subsequent steps in ICL repair are less well-defined. However, restoration of the reversed fork likely is required to deliver the 3’ end of the nascent leading strand up to the ICL, allowing translesion synthesis past the unhooked ICL adduct (“ICL remnant”) and completion of replication on one sister chromatid^10,11^. The two-ended DSB produced upon unhooking and fork reversal is can then repaired by homologous recombination using the fully-replicated sister chromatid as a template^12^.

**Figure 1.**
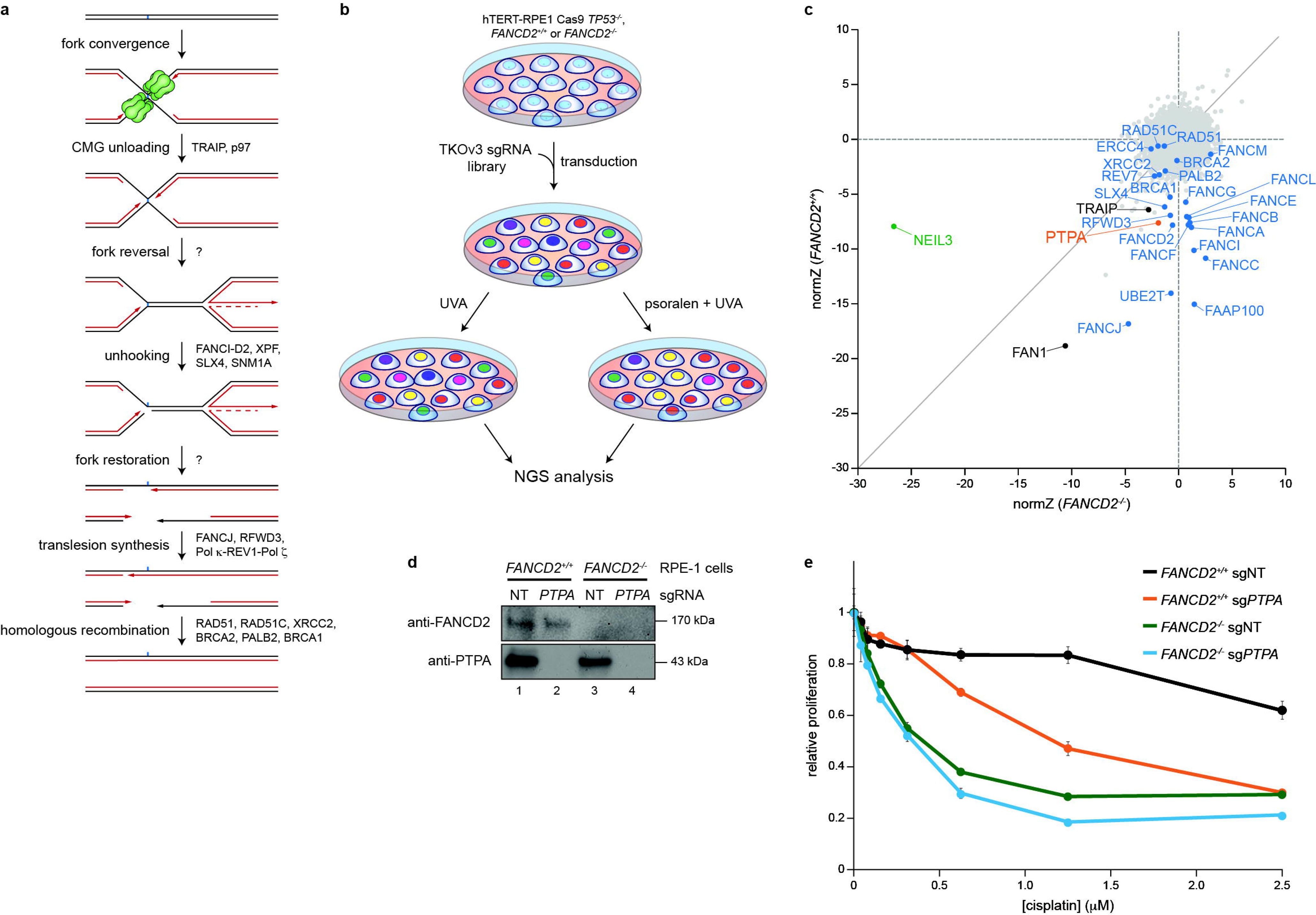
Genome-wide CRISPR dropout screens implicate PTPA in DNA interstrand cross-link repair. **a**, Model of replication-coupled ICL repair by the FA pathway. **b**, Schematic describing CRISPR dropout screens using human *FANCD2^+/+^* or *FANCD2^-/-^* RPE-1 cells. **c**, Biplot of normZ scores relating viability in psoralen and UVA treated *FANCD2^+/+^* (y-axis) and *FANCD2^-/-^* (x-axis) RPE-1 cells. **d**, Immunoblot demonstrating efficiency of PTPA depletion in *FANCD2^+/+^* and *FANCD2^-/-^* RPE-1 CRISPR knockout cell pools. Cells were transduced with lentivirus carrying non-targeting (NT) and *PTPA*-targeting sgRNAs. **e**, Relative RPE-1 cell proliferation after cisplatin treatment was measured using CellTiter-Glo reagent. *FANCD2^+/+^* and *FANCD2^-/-^* RPE-1 cells were transduced with sgRNAs as in **d**, and cell proliferation was measured after 5 days. Data represent the mean ± standard deviation from three technical replicates.

While current evidence indicates that replication fork reversal and restoration are integral steps in ICL repair by the FA pathway, little is known about the enzymes that execute these fork dynamics or how they are regulated. The enzyme that mediates fork reversal has not been identified, though the FANCM translocase is a likely candidate^13^. The factors responsible for fork restoration during ICL repair are also unknown, but this process may depend on one or more RECQ family translocases. The RECQ family members WRN and BLM can perform restoration of model reversed fork substrates *in vitro*^14,15^, and RECQ1 promotes fork restoration in response to camptothecin-induced replication stress^16^. Fork restoration during ICL repair may also depend on nuclease activities. For instance, the DNA2 nuclease-helicase and WRN helicase are known to act cooperatively to perform 5’ to 3’ resection of reversed fork regressed arms and promote replication restart following replication stress^17^. Currently, the mechanisms that regulate fork reversal and restoration are not well understood, though members of the phosphatidylinositol 3-kinase-related kinase (PIKK) family are thought to coordinate these processes. DNA-dependent protein kinase catalytic subunit (DNA-PKcs) and the ataxia-telangiectasia mutated (ATM) kinase promote fork reversal in response to replication stress^18–20^. Additionally, both ATM and ataxia-telangiectasia and Rad3-related (ATR) kinases phosphorylate resection enzymes, including WRN and MRE11, to maintain fork protection and ensure recovery from replication stress^21,22^. However, it is still unclear how these DNA damage checkpoint kinases are themselves activated and regulated in response to fork dynamics at sites of replication stress. Thus, the mechanisms governing replication fork remodeling in response to ICLs and other forms of replication stress remains a key unresolved question.

Here, we used genome-wide CRISPR screens and cell-free *Xenopus* egg extracts that faithfully model replication-coupled ICL repair to uncover a role for the ataxia and telangiectasia-mutated (ATM) checkpoint kinase in regulating reversed fork processing during ICL repair. We show that protein phosphatase inhibition disrupts ICL repair by the FA pathway by inducing aberrant resection of reversed fork intermediates and that this aberrant processing is suppressed by ATM inhibition. We further show that ATM is normally activated upon replication fork reversal and promotes resection of nascent DNA strands, which depends on DNA2, WRN, and EXO1. Our results support a model in which the regressed arm of a reversed fork functions as a one-ended DSB that activates ATM and stimulates DNA2- and EXO1-dependent resection of the regressed arm, thereby enabling fork restoration and completion of ICL repair.

## Results

### The PP2A phosphatase activator PTPA exhibits genetic epistasis with the FA pathway

To identify novel factors that participate in ICL repair by either the FA pathway or alternative repair pathways, we performed genome-wide chemogenomic CRISPR dropout screens with human hTERT-immortalized, p53-deficient RPE-1 cells (Fig. 1b and Supplementary Fig. 1)^23^. Cells were serially challenged with either UVA (control arm) or psoralen and UVA (PUVA; treatment arm), which induces the formation of psoralen-ICLs that are known to be repaired by both the FA and NEIL3 pathways^6,7,24^. Importantly, we performed screens using both FA pathway-proficient cells (*FANCD2^+/+^*) and isogenic FA pathway-deficient cells (*FANCD2^-/-^*). This approach enabled us to distinguish genes that function epistatically with the FA pathway (only conferring PUVA resistance in the FA pathway-proficient cells) from those that might function in alternative ICL repair pathways (conferring PUVA resistance in both FANCD2-proficient and - deficient cells). Validating our approach, we recovered 16 known *FANC* genes as ICL resistance genes in FANCD2-proficient cells (Fig. 1c). Importantly, except for *FANCJ*, none of these genes were recovered in FANCD2-deficient cells, consistent with roles for these genes in the same ICL repair pathway as *FANCD2*. Conversely, *NEIL3* and *FAN1*, both of which act independently of the FA pathway to promote ICL repair^24–27^, were recovered as ICL resistance genes in both FANCD2-proficient and -deficient cells.

In addition to known *FANC* genes, *PTPA* emerged as an ICL resistance gene specifically in FANCD2-proficient cells. *PTPA* encodes a factor that promotes PP2A phosphatase holoenzyme formation through activation of the PP2A catalytic subunit^28^, implying that PP2A promotes ICL repair by the FA pathway. Note that genes encoding PP2A subunits (e.g. PP2A-A and PP2A-C) are essential and would therefore be unlikely to emerge as hits in CRISPR dropout screens^29^. Consistently, another recent set of genome-wide screens also implicated *PTPA* in resistance to cisplatin, which induces ICLs that require the FA pathway for replication-coupled repair^23^. We therefore specifically tested whether *PTPA* contributes to ICL resistance. *PTPA* sgRNA knockout cell pools exhibited increased sensitivity to cisplatin (Fig. 1d,e), confirming a role for *PTPA* in cellular tolerance to cisplatin induced DNA damage. Overall, these results suggest that PP2A is required for ICL repair by the FA pathway.

### PP2A inhibition destabilizes reversed replication forks

To determine how PP2A phosphatase activity might regulate replication-coupled ICL repair, we replicated undamaged and ICL-containing plasmids in *Xenopus* egg extracts supplemented with the pan-phosphoprotein phosphatase (PPPase) inhibitor okadaic acid. Following replication in the presence of [α-^32^P]dCTP (which is incorporated into the nascent DNA strands), replication intermediates and products were deproteinized and resolved by native agarose gel electrophoresis (Fig. 2a). Since okadaic acid is known to block replication initiation by inhibiting incorporation of CDC45 into pre-replication complexes^30^, okadaic acid was added 5 minutes after initiating replication. Addition of okadaic acid did not appreciably affect replication of either an undamaged plasmid (pCtrl) or a plasmid containing an ICL formed by an abasic site (pICL^AP^), which is predominantly repaired by the NEIL3 pathway in egg extracts. In contrast, okadaic acid addition had a dramatic impact on replication of a plasmid containing a cisplatin-ICL (pICL^Pt^) that is repaired by the FA pathway^31^. In the absence of okadaic acid, pICL^Pt^ replication resulted in an initial accumulation of slowly-migrating figure 8 intermediates that correspond to the convergence of replication forks on either side of the ICL. These species were then converted into a topologically-distinct, faster-migrating figure 8 intermediate that forms upon unloading of stalled CMG helicases from the ICL. The fast figure 8 intermediates were in turn converted into species with intermediate mobilities that correspond to structures in which one of the replication forks abutting the ICL has undergone extensive reversal with resection of the nascent lagging strand^8^. Subsequent unhooking of the reversed fork intermediates through FANCI-FANCD2-dependent nucleolytic incisions resulted in the formation of sigma products and, eventually, fully-replicated, supercoiled repair products. Strikingly, addition of okadaic acid resulted in a rapid disappearance of reversed fork intermediates and a reduction in the accumulation of fully-replicated and repaired supercoiled products. This was accompanied by a transient, but more intense accumulation of sigma products and other faster migrating species (Fig. 2a, red arrowheads) corresponding to replication forks that have undergone cleavage of one or both parental DNA strands (Supplementary Fig. 2a,b). Okadaic acid also induced an earlier and more pronounced accumulation of well products, which may indicate the formation of end-joining intermediates and products following okadaic acid-induced fork cleavage^32^. Additionally, we observed the reappearance and persistence of species that co-migrate with fast figure 8 intermediates (Fig. 2a, black arrowhead), suggestive of premature reversed fork restoration prior to the occurrence of unhooking incisions. These observations suggest that PPPase activity promotes ICL repair by the FA pathway by ensuring proper processing of FA pathway repair intermediates. Treatment with the PP2A-selective inhibitor cytostatin^33^ similarly resulted in destabilization of pICL^Pt^ reversed fork intermediates and formation of fork cleavage products, well products, and putative cross-linked fork restoration products (Fig. 2b), indicating that PP2A, specifically, is required for efficient ICL repair by the FA pathway. Further, addition of excess recombinant human PP2A-AC subcomplex largely suppressed the effects of okadaic acid on pICL^Pt^ replication (Fig. 2c and Supplementary Fig. 2c,d).

**Figure 2.**
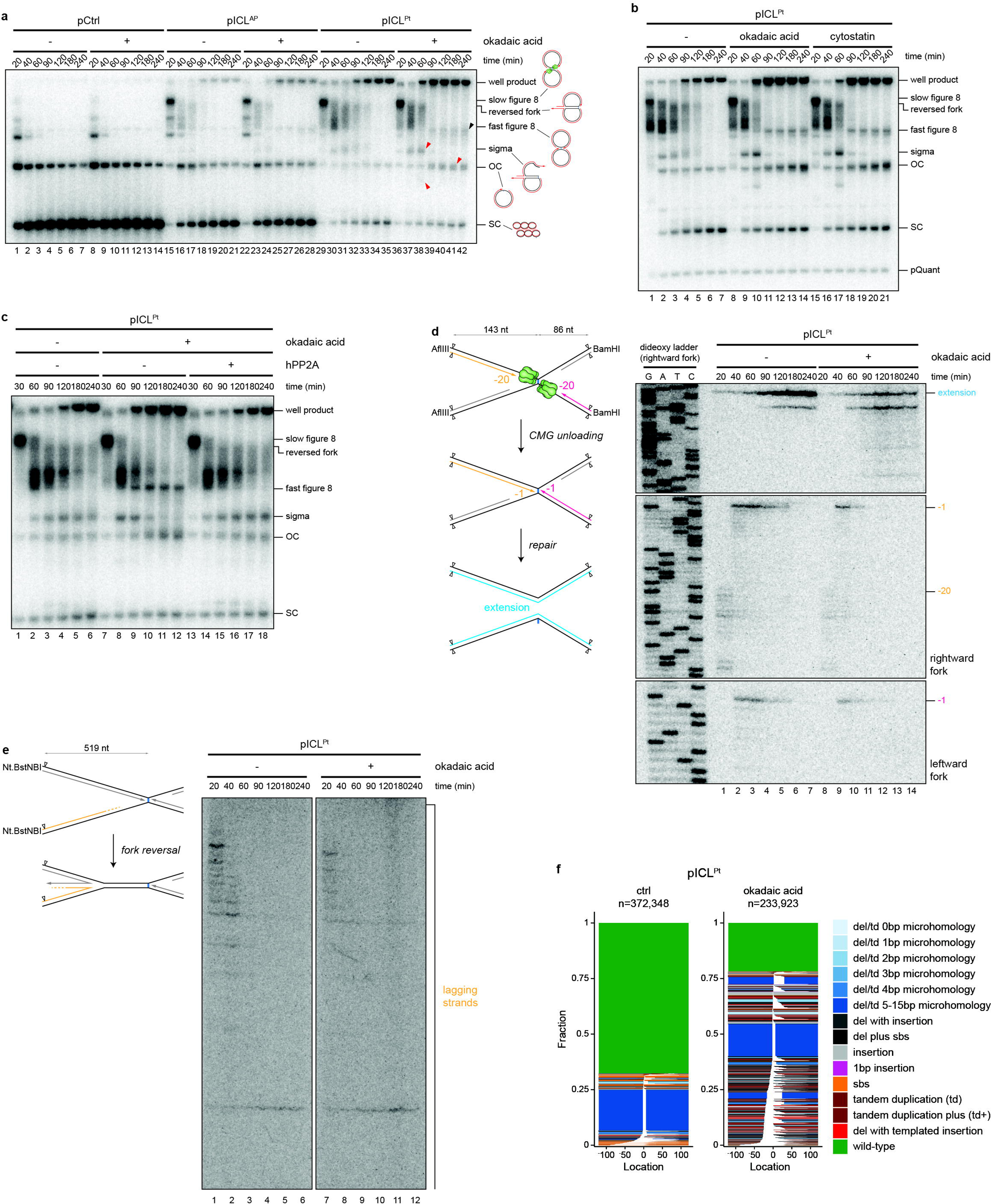
Phosphatase inhibition induces replication fork collapse during ICL repair. **a**, The indicated plasmids were replicated in egg extract supplemented with [α-^32^P]dCTP and the PPPase inhibitor okadaic acid, as indicated. Replication intermediates were recovered, deproteinized, separated on a native agarose gel, and visualized by autoradiography. Schematics of replication and repair intermediates that are resolved by native agarose gel electrophoresis are shown. SC, supercoiled; OC, open circular. Red arrowheads indicate fork cleavage products that accumulate during pICL^Pt^ replication in the presence of okadaic acid. Black arrowhead indicates putative fast figure 8 intermediates generated upon reversed fork restoration. **b**, pICL^Pt^ was replicated in egg extract supplemented with [α-^32^P]dCTP and okadaic acid or cytostatin, as indicated. Replication intermediates were analyzed as in **a**. **c**, pICL^Pt^ was replicated in egg extract supplemented with [α-^32^P]dCTP and okadaic acid and hPP2A-AC complex, as indicated. Replication intermediates were analyzed as in **a**. **d**, Left, schematic of nascent leading strands generated during ICL repair. AflIII cuts 143 nt to the left of the cisplatin-ICL. BamHI cuts 88 nt to the right of the cisplatin-ICL. Digestion with AflIII and BamHI generates characteristic −20 to −40 stall, −1 stall, insertion (“0”), and strand extension products. Right, pICL^Pt^ was replicated in egg extract supplemented with [α-^32^P]dCTP and okadaic acid, as indicated. Replication intermediates were isolated and digested with AflIII and BamHI, and nascent DNA strands were resolved by denaturing polyacrylamide gel electrophoresis. Top, middle, and bottom panels show sections of the same gels to visualize extension, rightward leading strands, and leftward leading strands, respectively. **e**, Left, schematic of nascent lagging strands generated during ICL repair. Nt.BstNBI nicks the rightward nascent lagging strand 519 nt to the left of the cisplatin-ICL. Right, pICL^Pt^ was replicated as in **d**. Replication intermediates were isolated and digested with Nt.BstNBI, and nascent DNA strands were resolved by denaturing polyacrylamide gel electrophoresis. **f**, pICL^Pt^ was replicated in egg extract supplemented with okadaic acid, and repair products were PCR amplified and sequenced. Raw paired-end sequencing reads were merged using SIQ, and data were visualized with SIQPlotteR. Color-coded repair outcomes are depicted as tornado plots. White spaces depict deletions. The positions of variations are indicated relative to the cisplatin-ICL site (“0”). n indicates the number of mapped sequencing reads obtained for each condition.

To better understand how okadaic acid disrupts ICL repair by the FA pathway, we monitored the stability and maturation of the nascent DNA strands during ICL repair. pICL^Pt^ was replicated in egg extract supplemented with [α-^32^P]dCTP and okadaic acid, as indicated, and recovered replication intermediates were digested with restriction enzymes that allow the nascent DNA leading strands to be resolved on a denaturing sequencing gel (Fig. 2d). In the absence of okadaic acid, nascent leading strands initially stalled 20 to 40 nucleotides (nt) upstream of the ICL (−20 to −40 stall) due to the footprint of CMG. Upon CMG unloading, the nascent strands were extended up to 1 nt upstream of the ICL (−1 stall). ICL unhooking and TLS past the resulting ICL remnant converted the nascent leading strands into full length extension products. In the presence of okadaic acid, nascent leading strands first stalled at −20 to −40, before being extended to the −1 position. However, okadaic acid treatment induced a rapid disappearance of nascent leading strand stall products. This loss of leading strand stall products was accompanied by both a decrease in formation of expected full-length nascent strand extension products and an increase in nascent strand extension products that were predominantly shorter than the expected product, suggesting that excessive okadaic acid-induced processing of replication intermediates leads to fork collapse and formation of deletion-prone end-joining products. We also specifically examined the fate of nascent lagging strands using a nicking enzyme that liberates the lagging strand from one fork. Okadaic acid treatment accelerated the disappearance of nascent lagging strands (Fig. 2e; compare lanes 1 and 2 with lanes 7 and 8), consistent with accelerated 5’ to 3’ resection of reversed fork intermediates. Sequencing of pICL^Pt^ replication products recovered from untreated or okadaic acid-treated extracts confirmed that okadaic acid treatment caused a reduction in the fraction of sequencing reads corresponding to error-free repair products and a substantial increase in the fraction of reads with deletions spanning the ICL (Fig. 2f). Taken together, our results support a model in which PP2A limits resection of repair intermediates, which can lead to fork collapse and error-prone end-joining.

### ATM promotes reversed fork processing during ICL repair

PP2A targets thousands of phosphosites across the human proteome^34^, making the identification of specific dephosphorylation events that regulate ICL repair a formidable challenge. However, we reasoned that PP2A activity would likely counteract one or more protein kinases that are activated during ICL repair. Accordingly, inhibition of relevant kinase activities should prevent excessive resection of reversed fork intermediates during ICL repair in phosphatase-inhibited egg extract. We therefore tested whether chemical inhibition of CDKs or the apical DNA damage response kinases ATR or ATM suppresses the effects of okadaic acid on replication intermediate processing during pICL^Pt^ replication in egg extract. Pan-CDK inhibition with flavopiridol did not noticeably affect the formation and resolution of pICL^Pt^ repair intermediates and did not suppress okadaic acid-induced destabilization of reversed fork intermediates (Supplementary Fig. 3a,b). On its own, addition of the ATR inhibitor AZD6738 caused a subtle accumulation of reversed fork intermediates (Supplementary Fig. 3c), consistent with a role for ATR in stimulating DSB end resection^35^. However, ATR inhibition did not substantially increase the stability of reversed fork intermediates or suppress the accumulation of putative fork cleavage products in okadaic acid-treated extract (Supplementary Fig. 3c). In contrast, addition of the ATM inhibitor KU-55933 during pICL^Pt^ replication resulted in a strong accumulation of reversed fork intermediates (Fig. 3a). KU-55933 addition also increased the stability of reversed fork intermediates and delayed the formation of fork cleavage products during pICL^Pt^ replication in okadaic acid-treated extract, suggesting that PP2A counteracts ATM to regulate processing of reversed fork intermediates during ICL repair. We observed similar stabilization of reversed forks—both in the presence and absence of okadaic acid—in extracts supplemented with another ATM inhibitor (AZD0156) as well as with caffeine, an inhibitor of both ATM and ATR (Supplementary Fig. 3d,e). Consistent with the idea that PP2A counteracts ATM activation, okadaic acid treatment during pICL^Pt^ replication induced increased phosphorylation of ATM at S1981, a canonical marker of ATM activation, as well as phosphorylation of the ATM target proteins MRE11 and CHK2 (Fig. 3b). This phosphorylation of ATM, MRE11, and CHK2 was attenuated by addition of ATM inhibitor. Combined, our results indicate that PP2A prevents hyperactivation of ATM during ICL repair. This conclusion aligns with previous work demonstrating that PP2A interacts with ATM and suppresses ATM activation in the absence of DNA damage^36^.

**Figure 3.**
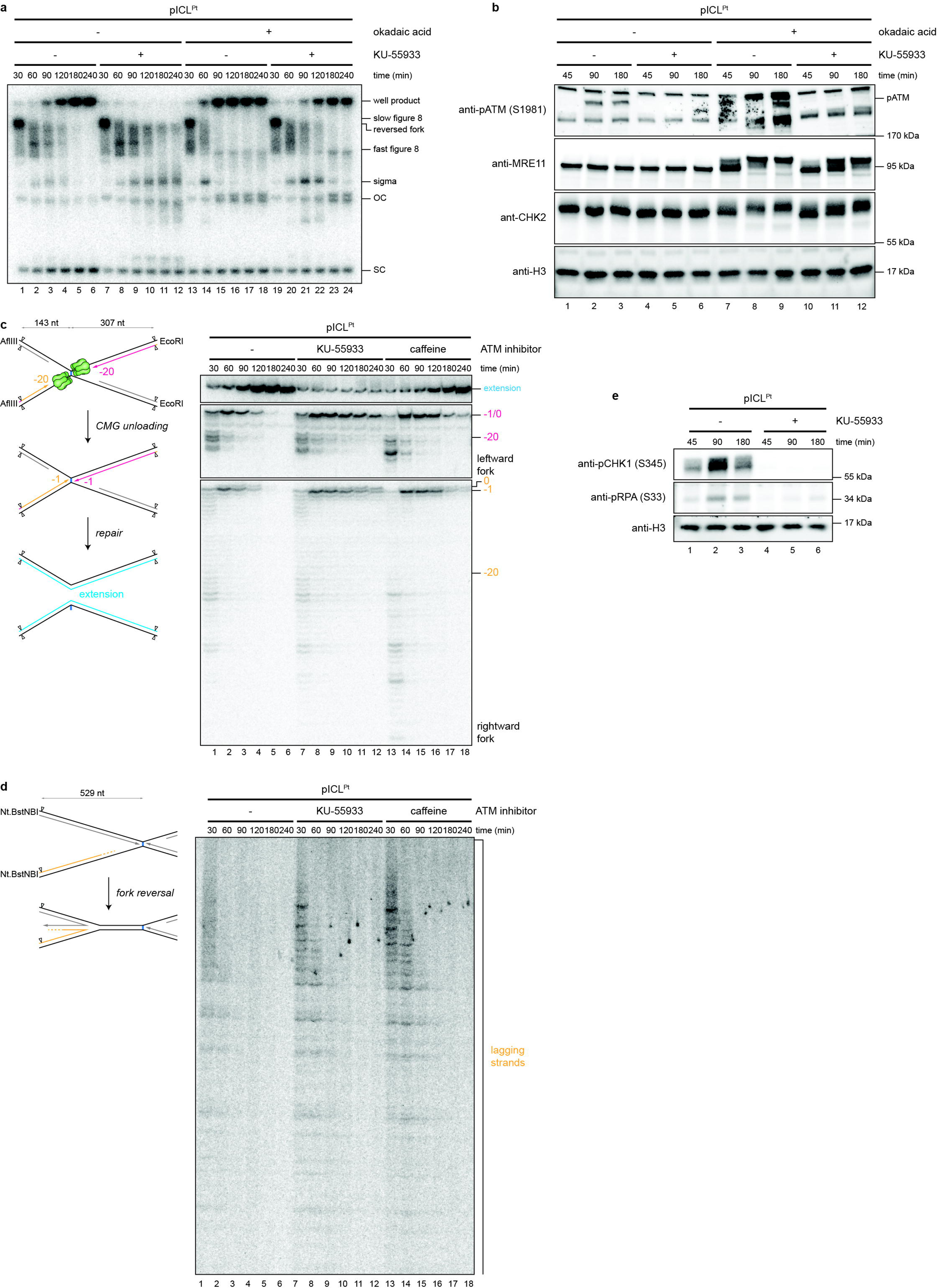
ATM promotes reversed fork processing during ICL repair. **a**, pICL^Pt^ was replicated in egg extract supplemented with [α-^32^P]dCTP and okadaic acid and the ATM inhibitor KU-55933, as indicated. Replication intermediates were analyzed as in Fig. 2a. **b**, pICL^Pt^ was replicated in egg extract supplemented with okadaic acid and KU-55933, as indicated. Replication reactions were separated by SDS–PAGE and blotted for phospho-ATM (S1981) and H3 (loading control). **c**, Nascent DNA leading strands from pICL^Pt^ replication reactions supplemented with [α-^32^P]dCTP and KU-55933 or caffeine were digested with AflIII and EcoRI and analyzed as in Fig. 2d. AflIII cuts 143 nt to the left of the cisplatin-ICL. EcoRI cuts 307 nt to the right of the cisplatin-ICL. **d**, Nascent DNA lagging strands from pICL^Pt^ replication reactions supplemented with [α-^32^P]dCTP and KU-55933 or caffeine were digested with Nt.BstNBI and analyzed as in Fig. 2e. **e**, pICL^Pt^ was replicated in egg extract supplemented with KU-55933, as indicated. Replication reactions were blotted for phospho-CHK1 (S345), phospho-RPA32 (S33) and H3 (loading control) as in **b**.

We then determined how ATM inhibition influences the processing and maturation of nascent DNA strands. ATM inhibition with either KU-55933 or caffeine caused a strong accumulation of nascent leading strand −1 and 0 stall products and a corresponding reduction in full length extension products (Fig. 3c). ATM inhibition also resulted in stabilization of nascent lagging strands (Fig. 3d). Combined, these results suggest that ATM may stimulate resection of reversed fork intermediates and thereby facilitate extension of the nascent leading strand past an unhooked ICL. Alternatively, stabilization of leading and lagging strands in ATM-inhibited extract could stem from defective homologous recombination and a failure to complete repair of DSBs generated during ICL repair. Indeed, in addition to stabilizing reversed fork intermediates, ATM inhibition also stabilized linear plasmids believed to form upon fork restoration and suppressed the formation of well products thought to correspond to HR intermediates (Fig. 3a). To distinguish whether ATM promotes resection during ICL repair, we examined the extent of CHK1 and RPA phosphorylation during pICL^Pt^ replication in untreated and ATM-inhibited extract. CHK1 and RPA are phosphorylated by ATR upon accumulation of single stranded (ss)DNA and serve as markers of DNA resection^37^. ATM inhibition strongly suppressed the accumulation of phosphorylated forms of CHK1 and RPA (Fig. 3e), indicating that ATM inhibition suppresses the accumulation of ssDNA during ICL repair. In aggregate, our data indicate that ATM promotes resection of reversed fork intermediates during ICL repair. This conclusion is consistent with ATM’s established role in promoting resection of canonical DSB substrates^38^.

### ATM is activated concomitantly with fork reversal

We next tested whether ATM becomes activated by a specific DNA structure during ICL repair. pCtrl and pICL^Pt^ were replicated in egg extracts, and ATM activation was monitored by immunoblotting for ATM phosphorylation at S1981. Whereas ATM phosphorylation was undetectable during mock replication in the absence of plasmid and during replication of undamaged pCtrl, phosphorylated ATM accumulated in a time-dependent manner during pICL^Pt^ replication (Fig. 4a), indicating that ATM activation is damage dependent. During ICL repair by the FA pathway, fork reversal produces a regressed DNA arm that is equivalent to a one-ended DSB that we suspected might trigger ATM activation. Consistent with this hypothesis, ATM phosphorylation was not activated during pICL^Pt^ replication in extract supplemented with p97 inhibitor (Fig. 4a and Supplementary Fig. 4a), which blocks CMG loading and subsequent fork reversal. A DSB is also generated through FANCI- and FANCD2-dependent incisions that unhook the ICL^11^. We therefore examined the extent of ATM activation during pICL^Pt^ replication in FANCD2-depleted extract. FANCD2-depletion led to a pronounced reduction in the amount of sigma intermediate incision products (Supplementary Fig. 4b,c) but did not diminish the extent of ATM phosphorylation (Fig. 4b). We conclude that the reversed fork structure generated during ICL repair is sufficient to activate ATM and processing of the regressed arm.

**Figure 4.**
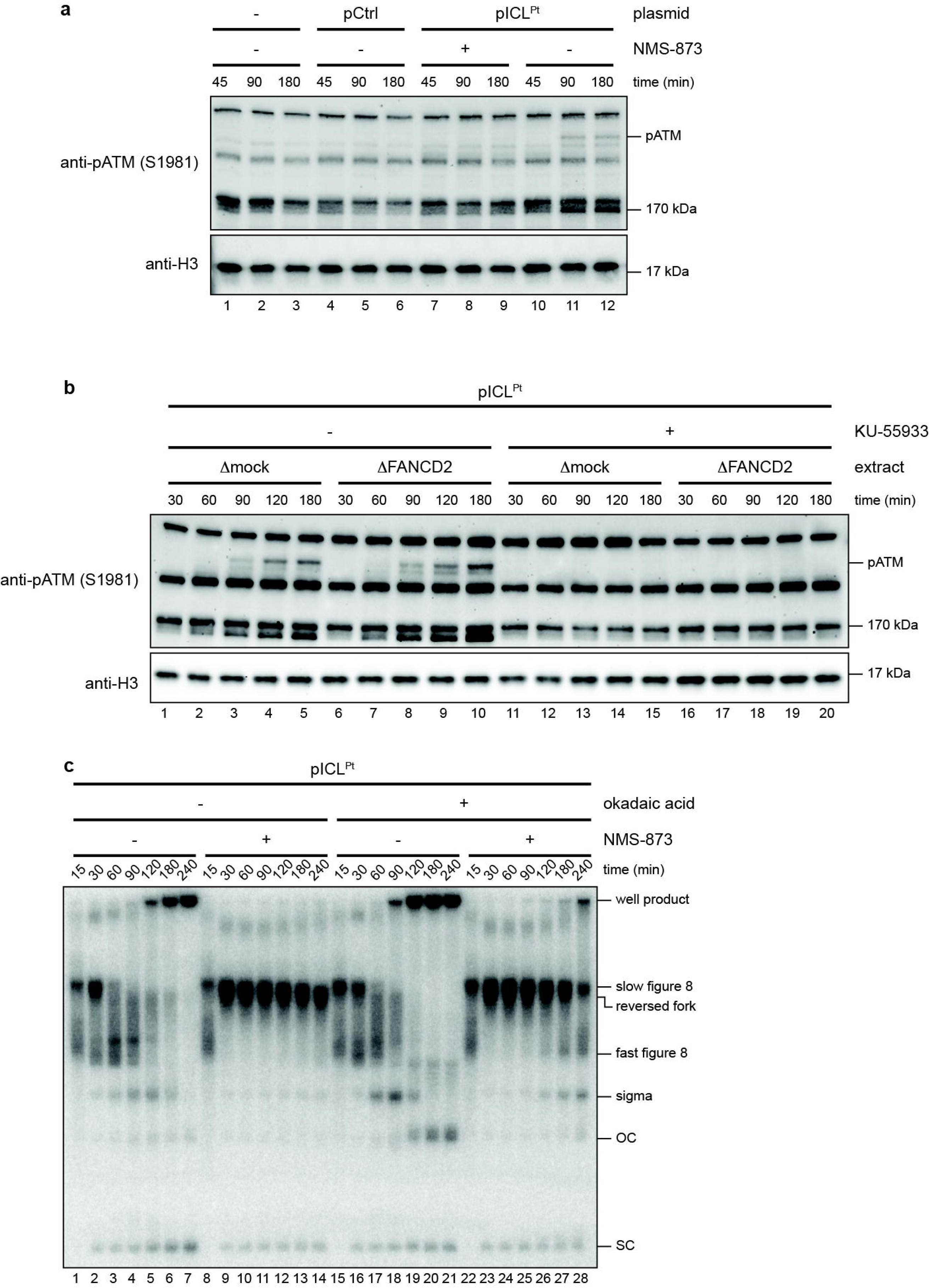
ATM activation during ICL repair requires CMG unloading. **a**, Plasmids were replicated in egg extract supplemented with okadaic acid and the p97 inhibitor NMS-873, as indicated. Replication reactions were blotted for phospho-ATM (S1981) and H3 as in Fig. 3b. **b**, pICL^Pt^ was replicated in mock- or FANCD2-depleted egg extract supplemented with KU-55933, as indicated. Replication reactions were blotted for phospho-ATM (S1981) and H3 as in Fig. 3b. **c**, pICL^Pt^ was replicated in egg extract supplemented with [α-^32^P]dCTP and okadaic acid and the p97 inhibitor NMS-873, as indicated. Replication intermediates were analyzed as in Fig. 2a.

We also determined whether ATM hyperactivation can induce processing of replication intermediates other than reversed fork structures. pICL^Pt^ was replicated in egg extract that was supplemented with okadaic acid and p97 inhibitor, as indicated, and replication intermediates were resolved on a native agarose gel (Fig. 4c). As expected, in the absence of okadaic acid, p97 inhibitor blocked the formation of reversed fork intermediates and caused a persistence of slow figure 8 intermediates that form when forks converge at the ICL. Despite high levels of basal ATM activation (Fig. 4a), slow figure 8 intermediates also persisted in extract supplemented with both okadaic acid and p97 inhibitor, indicating that these structures are resistant to ATM-induced processing (Fig. 4c). The slow figure intermediates remained stable in okadaic acid-treated extracts in which ATM activation was further stimulated by addition of linear plasmid DNA (Supplementary Fig. 4d,e). These data suggest that ATM-dependent nucleolytic processing is specifically targeted to reversed fork intermediates.

### DNA2, WRN, and EXO1 mediate reversed fork resection during ICL repair

To identify potential effectors of ATM-dependent reversed fork resection, we profiled the phosphoproteome during pICL^Pt^ replication in untreated and okadaic acid-treated extracts. Okadaic acid treatment resulted in a significant, at least 2-fold increase in abundance for ∼8.7% of phosphopeptides detected (Fig. 5a and Supplementary Fig. 5a,b). A number of the phosphopeptides that were significantly enriched in okadaic acid-treated extract corresponded to the nucleases DNA2 and EXO1 as well as the DNA2-associated helicases BLM and WRN. Since DNA2, WRN, and EXO1 have been previously implicated in reversed fork processing^17,39^, we tested whether they mediate resection during ICL repair. Addition of the DNA2 inhibitor C5^40^, the EXO1 inhibitor C200^41^, or the WRN inhibitor HRO761^42^ to extracts resulted in a strong accumulation of reversed fork intermediates and a decrease in the amount of fully-replicated supercoiled products, mirroring the effects of ATM inhibition and suggesting that DNA2, WRN, and EXO1 process the reversed fork intermediates and enable completion of ICL repair (Fig. 5b). Similar stabilization of reversed fork intermediates generated during ICL repair was also observed with the EXO1 inhibitor F684^41^ (Supplementary Fig. 5c). Inhibition of DNA2, WRN, or EXO1 also stabilized reversed fork intermediates and delayed the accumulation of putative fork collapse products in okadaic acid-treated extract (Fig. 5b), implying that these nucleases and helicases drive reversed fork over-resection in the absence of PP2A activity.

**Figure 5.**
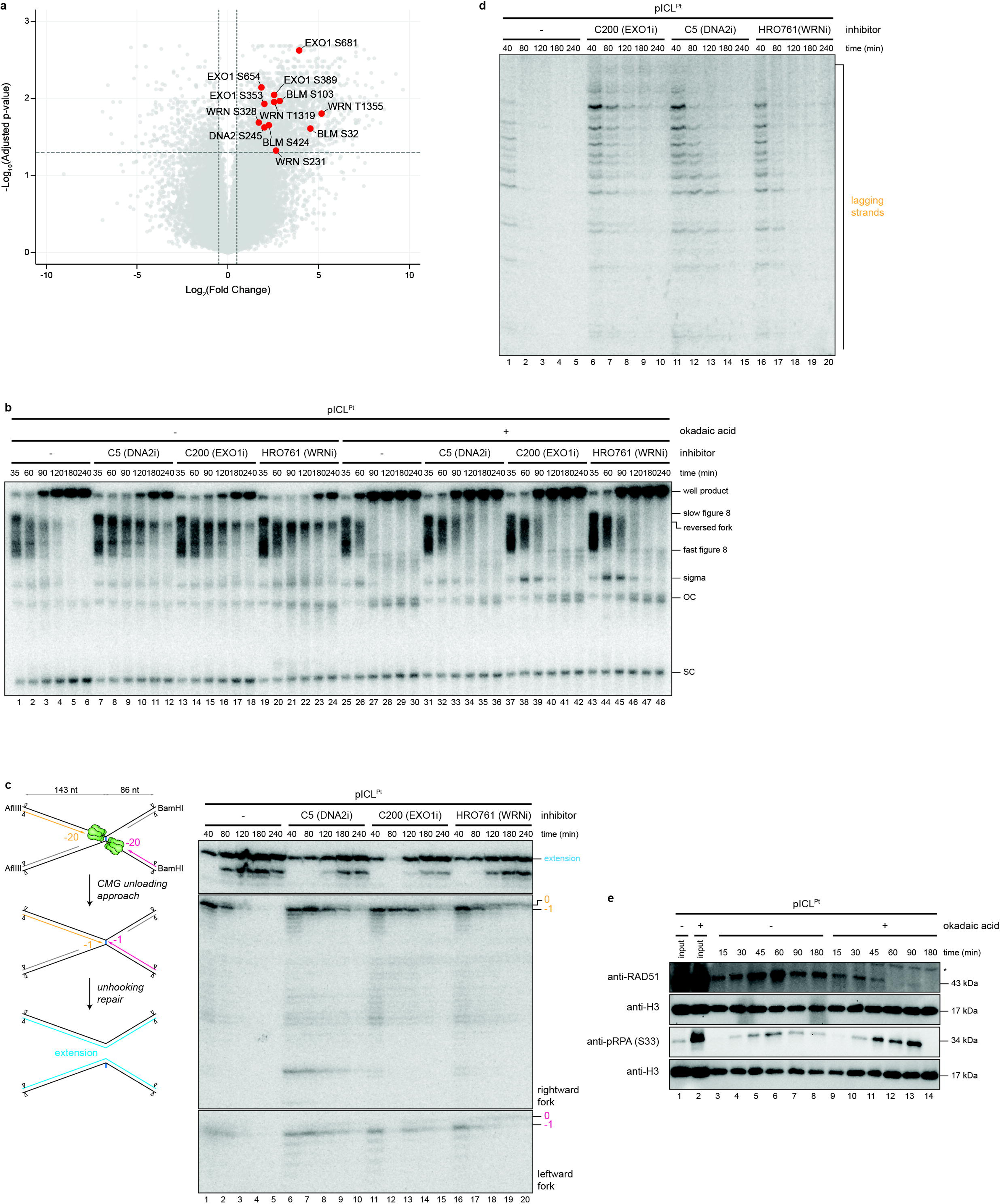
EXO1 and DNA2 mediate processing of reversed fork intermediates during ICL repair. **a**, Volcano plots comparing the abundance of phosphopeptides 90 minutes after initiating pICL^Pt^ replication in untreated and okadaic acid-supplemented egg extract. The estimated fold change (FC) in normalized phosphopeptide intensity between the samples is plotted against the false discovery rate-(FDR-)adjusted p-value. Fold changes (FCs) and p-values were computed using linear mixed-effects models implemented in the MSstatsPTM package^63^. p-values were adjusted for multiple hypothesis testing using the Benjamini-Hochberg procedure^64^. Labeled proteins of interest meet significance criteria (|Log_2_FC| > 0.5 and p < 0.05). **b**, pICL^Pt^ was replicated in egg extract supplemented with [α-^32^P]dCTP and okadaic acid and the DNA2 inhibitor C5, the EXO1 inhibitor C200, or the WRN inhibitor HRO761, as indicated. Replication intermediates were analyzed as in Fig. 2a. **c**, Nascent DNA leading strands from pICL^Pt^ replication reactions supplemented with [α-^32^P]dCTP and C5, C200, or HRO761 were digested with AflIII and BamHI and analyzed as in Fig. 2d. **d**, Nascent DNA lagging strands from pICL^Pt^ replication reactions supplemented with [α-^32^P]dCTP and C200, C5, or HRO761 were digested with Nt.BstNBI and analyzed as in Fig. 2e. **e**, pICL^Pt^ was replicated in egg extract supplemented with okadaic acid, and recovered chromatin was blotted for RAD51, phospho-RPA32 (S33), and H3.

We next examined the effects of DNA2, WRN, and EXO1 inhibition on nascent strand maturation. Like ATM inhibition, inhibition of DNA2, WRN, or EXO1 caused a strong persistence of −1 and 0 leading strand stall products and a corresponding reduction in the amount of full-length extension products (Fig. 5c), suggesting that EXO1, DNA2, and WRN might be required for reversed fork restoration and subsequent bypass of the unhooked ICL remnant. Inhibition of DNA2, WRN, or EXO1 also caused a strong persistence of nascent lagging strands, indicating that these nucleases promote 5’ to 3’ resection of reversed forks (Fig. 5d). Interestingly, EXO1 inhibition resulted in a more pronounced stabilization of the longest lagging strands with 5’ ends nearest to the end of the regressed arm of a reversed fork structure. In contrast, inhibition of DNA2 or WRN resulted in more pronounced stabilization of shorter lagging strands corresponding to recessed 5’ ends on the regressed arm of a reversed fork. These data suggest a model in which EXO1 initiates 5’ to 3’ resection at a reversed fork, after which DNA2 acts with WRN to further resect 5’ recessed ends.

Reversed fork processing by DNA2-WRN and EXO1 is suppressed by RAD51-dependent fork protection^39,43^. To determine whether PPPase inhibition results in defective fork protection, we replicated pICL^Pt^ in untreated or okadaic acid-treated extract, recovered chromatin, and monitored RAD51 association by immunoblotting. Okadaic acid treatment resulted in a reduction in chromatin-associated RAD51 and a corresponding accumulation of phosphorylated RPA (Fig. 5e). This result suggests that aberrant DNA2-, WRN-, and EXO1-dependent processing of reversed forks during ICL repair might be at least partially explained by a failure to efficiently load protective RAD51 filaments onto the reversed fork intermediate. However, addition of BRC peptide that attenuates loading of RAD51 onto chromatin during ICL repair in egg extracts^12^ did not induce destabilization of reversed fork intermediates, as observed with okadaic acid treatment (Fig. S5d). Compromised RAD51-dependent fork protection therefore does not appear to be sufficient to induce degradation of reversed fork intermediates during ICL repair.

## Discussion

Although current data indicate that replication fork reversal and restoration are integral steps in ICL repair by the FA pathway, little is known about which factors mediate these fork dynamics or how they are regulated. Here, we used chemogenomic screens and *Xenopus* egg extracts to uncover a phosphoregulatory circuit involving ATM that contributes to reversed fork processing during ICL repair. A CRISPR dropout screen implicated the PP2A activator PTPA in ICL repair by the FA pathway. We then determined that inhibition of protein phosphatase activity results in hyperactivation of ATM and is associated with destabilization of reversed fork intermediates, fork collapse, and formation of deletion-prone end-joining products. Our data indicate that ATM is activated concomitantly with fork reversal and that this activation is required for 5’ to 3’ resection of nascent lagging strands (Fig. 6*i*). These observations indicate that ATM directs resection of reversed forks in a manner that is counteracted by phosphatase activity. We further implicate the long-range resection enzymes DNA2-WRN and EXO1 in mediating this resection, implying that these factors are directly or indirectly activated by ATM at a reversed fork. Thus, our results indicate that the reversed fork formed during ICL repair is recognized as a DSB and initiates a checkpoint signaling cascade similar to that triggered by a canonical, damage-induced DSB. Given that replication fork reversal is a frequent response to diverse replication stresses^44^, it seems likely that ATM activation regulates reversed fork processing outside the context of ICL repair.

**Figure 6.**
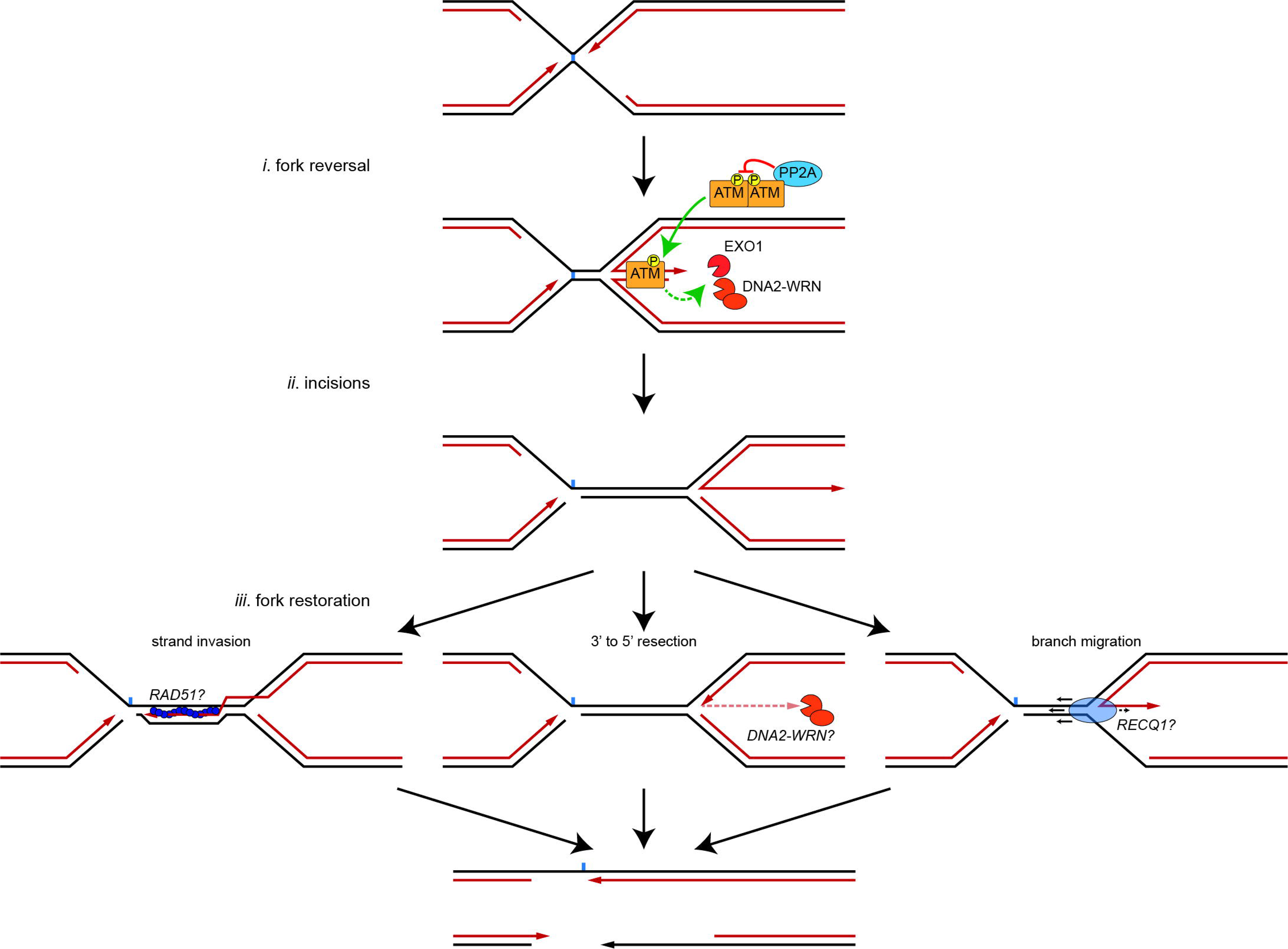
Model for regulation of reversed fork processing by ATM. See main text for details. Fork reversal (*i*) generates a regressed arm that activates ATM. ATM activation directly or indirectly stimulates nascent strand resection by EXO1 and DNA2-WRN. PP2A suppresses ATM activation to constrain resection. Following ICL unhooking by the FA pathway (*ii*), fork restoration (*iii*) enables completion of replication and repair. Potential mechanisms by which resection could promote fork restoration are shown.

Undoubtedly, ATM and phosphatase inhibition have pleiotropic effects on ICL repair. Consequently, the mechanisms by which phosphoregulation contributes to ICL repair remain to be determined. ATM is known to phosphorylate EXO1 and to physically interact with DNA2^45,46^. ATM also phosphorylates WRN and BLM helicases, both of which stimulate DNA2-dependent resection and have been implicated in the processing of reversed forks^17,47–51^. Thus, ATM may directly activate DNA2, WRN, and/or EXO1 to stimulate resection at reversed forks. Indirect mechanisms may also contribute to resection. PP2A interacts with BRCA2 to promote RAD51 filament formation during homologous recombination^52^. Inhibition of PP2A could therefore compromise RAD51-dependent protection of reversed replication forks, indirectly leading to excessive resection by EXO1 and DNA2-WRN. While both our data and previous work^36^ indicate that PP2A antagonizes ATM activation by dephosphorylating ATM itself, it seems likely that other phosphatases limit reversed fork resection by directly dephosphorylating enzymes that are activated through ATM-dependent signaling. The RIF1-PP1 complex, for example, is known to suppress DNA2-dependent fork degradation^53,54^. Additionally, ATM can promote fork reversal by stimulating recruitment of the SMARCAL1 translocase to stressed replication forks^18^. Thus, ATM appears to regulate multiple facets of fork reversal dynamics. Future investigations will be necessary to determine how ATM and relevant phosphorylation-dephosphorylation events balance fork reversal and processing to ensure productive recovery from replication stress.

Another key outstanding question concerns how ATM-stimulated reversed fork resection might promote fork restoration and the completion of ICL repair. We can envision multiple possibilities. 5’ to 3’ resection of the nascent lagging strand could allow RAD51 to load onto the resulting 3’ overhang and mediate invasion of the parental DNA duplex, generating a joint molecule structure with the nascent leading strand poised for extension past the unhooked ICL remnant (Fig. 6*iii*, left). While this idea may be consistent with available data, evidence for the formation of strand invasion intermediates predicted by this model is currently lacking^8^. 5’ to 3’ resection could also act in concert with a 3’ to 5’ exonuclease or 3’ flap endonuclease activity that degrades the 3’ overhang, eventually regenerating a replication fork with a free 3’ end that that can be extended through the parental DNA duplex and past the ICL remnant (Fig. 6*iii*, center). Interestingly, our data indicate that nascent leading strand −1 stall products are highly unstable in extract treated with phosphatase inhibitor, though it is unclear whether these nascent leading strands are processed though end-joining or nucleolytic degradation. Nonetheless, it is worth noting that DNA2, which our data implicate in reversed fork processing during ICL repair, has robust 3’ to 5’ nuclease activity that can, in some contexts, mediate nascent leading strand degradation at a reversed fork^55,56^. Resection of the regressed arm might also cooperate with branch migration to facilitate fork restoration by a DNA translocase, such as a member of the RECQ family (Fig. 6*iii*, right). If targeted to both strands of the regressed arm, resection would reduce the distance of branch migration required before a replication fork is restored. It is also possible that by converting the regressed arm into a ssDNA 3’ flap, resection imparts a polarity at the reversed fork four-way junction that biases branch migration in favor of fork restoration rather than further reversal. ATM-dependent reversed fork processing could also indirectly favor fork restoration by antagonizing toxic non-homologous end-joining of reversed fork regressed arms^57,58^. Alternatively, ATM-stimulated end resection could have a neutral or even deleterious effect on fork restoration. In this case, resection might reflect specious ATM activation at a reversed fork that is normally suppressed by phosphatases.

Our results suggest a mechanistic basis for understanding previously reported genetic interactions between DNA2 and the FA pathway. Depletion of both DNA2 and EXO1 induces cellular hypersensitivity to cisplatin, implying that resection is required during ICL repair^59^. Counterintuitively, DNA2 depletion suppresses ICL sensitivity in FANCD2-deficient cells, indicating that DNA2-dependent resection of ICL repair intermediates can corrupt ICL repair when the FANCD2 function is compromised^60^. We propose that DNA2-deficiency can suppress defects in the FA pathway by extending the window during which unhooking can occur before nonproductive fork restoration or excessive fork resection renders the ICL refractory to structure specific nucleases. This model suggests that manipulating the balance between fork reversal and fork restoration could have therapeutic benefit in FA patients. Low doses of ATM inhibitors could stabilize reversed fork intermediates to facilitate ICL unhooking by a compromised FA pathway, though FA-deficient cells are profoundly sensitive to ATM inhibition due to an inability to activate homologous recombination and suppress toxic non-homologous end-joining^61^. However, it may be possible to dampen reversed fork resection and rescue ICL repair using small molecule activators of PP2A (SMAPs)^62^.

---

**Supplementary Figure 1.**
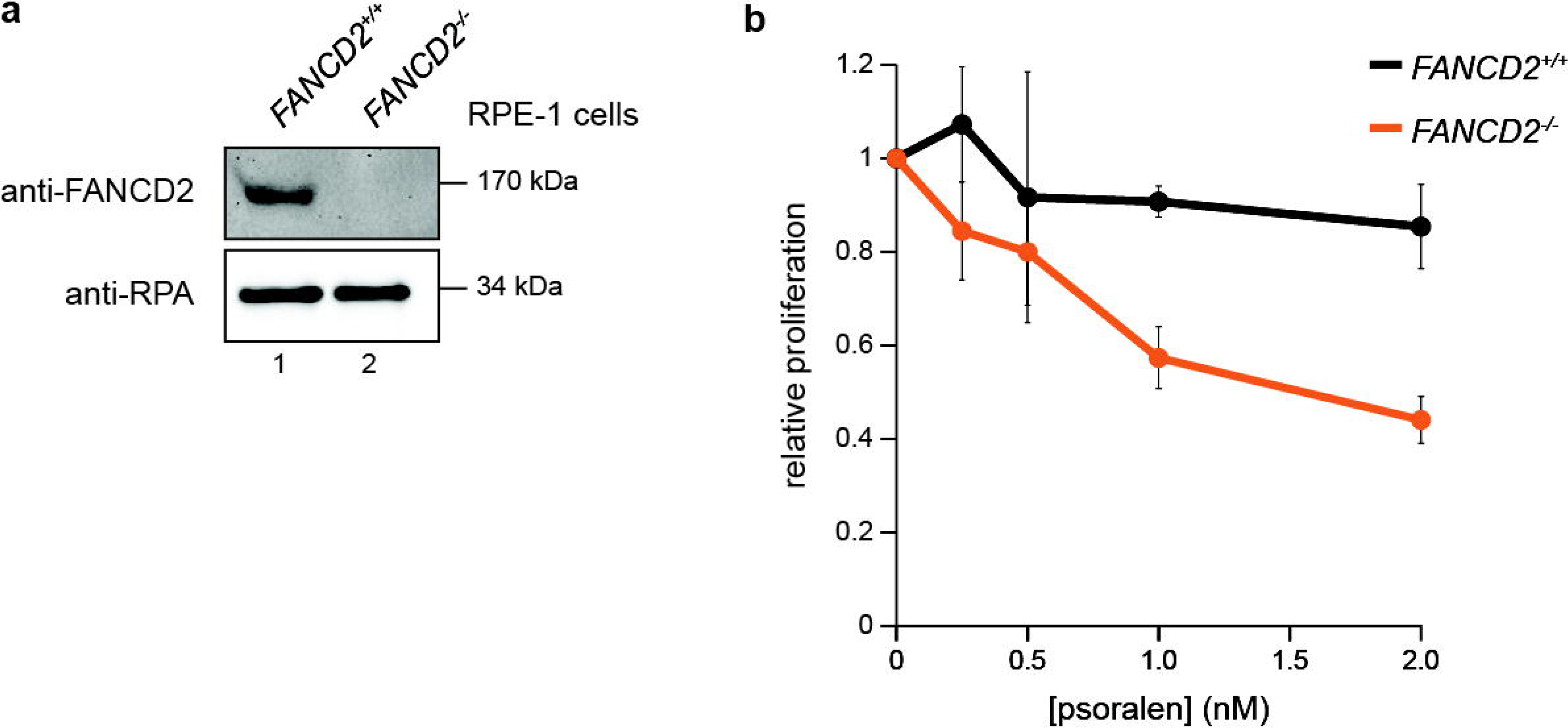
Validation of RPE-1 cells used in CRISPR dropout screen. **a**, Lysates from RPE-1 cells used for the chemogenomic screens described in Fig. 1b,c were blotted for FANCD2 and RPA32 (loading control). **b**, Relative *FANCD2^+/+^* and *FANCD2^-/-^* RPE-1 cell proliferation after psoralen + UVA treatment was measured using CellTiter-Glo reagent. Cell proliferation was measured after 5 days. Data represent the mean ± standard deviation from two independent experiments.

**Supplementary Figure 2.**
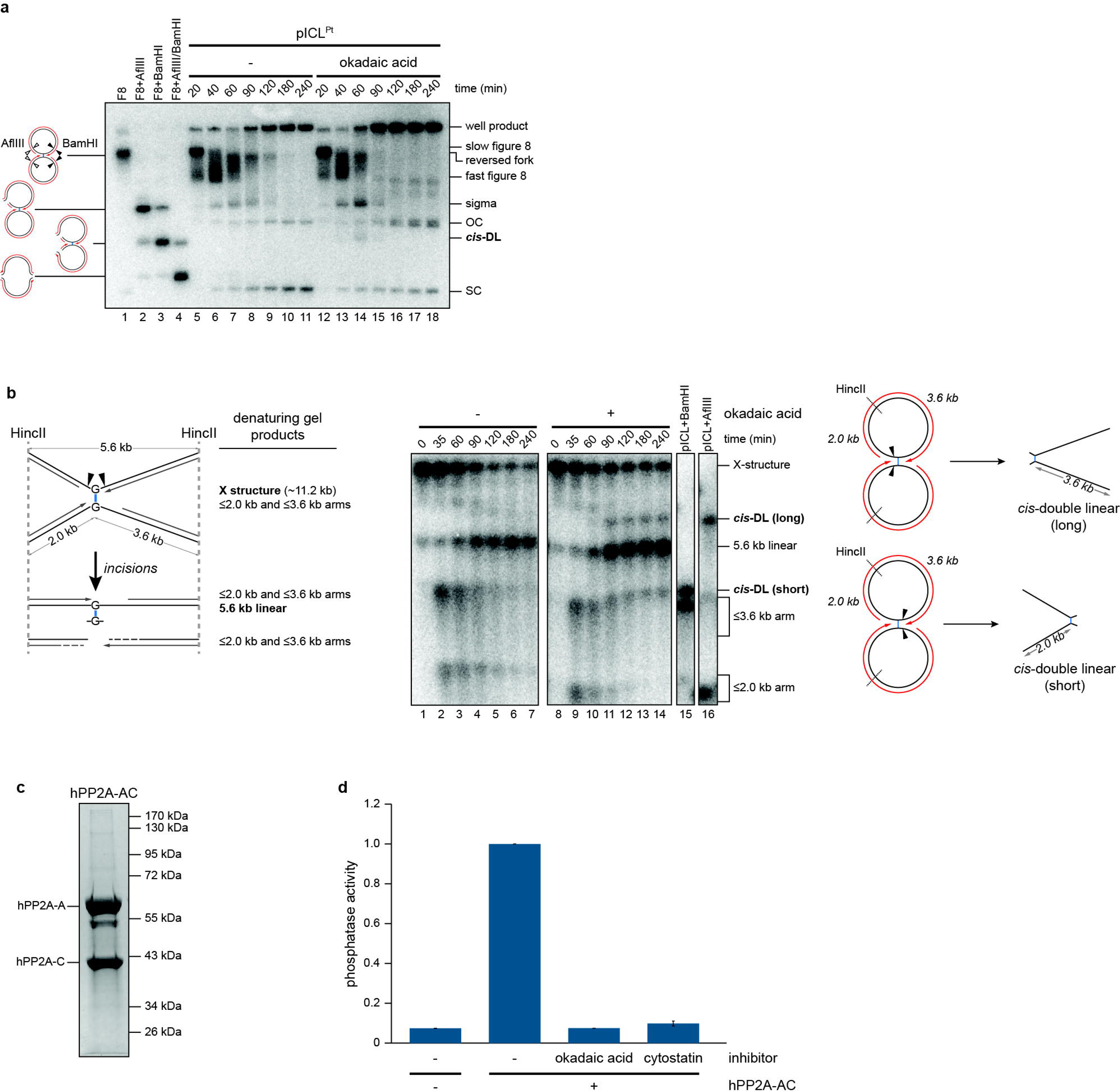
Characterization of okadaic acid-induced fork collapse products. **a**, pICL^Pt^ was replicated in egg extract supplemented with [α-^32^P]dCTP and okadaic acid, as indicated. Replication intermediates were resolved on a native agarose gel as in Fig. 2a. The identities of standards generated by digesting cross-linked figure 8 replication intermediates with AflIII (white arrowheads) and/or BamHI (black arrowheads) are shown to the left of the gel. A single cut by either AflIII or BamHI produces a species that migrates like a sigma intermediate, while two cuts produce a species that migrates similarly to a 10 kilobase pair linear DNA (*cis*-double linear). Digestion with both AflIII and BamHI produces a ∼5.6 kilobase pair linear. Note that an okadaic acid induced putative cleavage product comigrates with the *cis*-double linear cleavage standard, suggesting this species is produced upon cleavage of both parental DNA strands at one fork. **b**, Left, schematic of species generated during ICL repair. Digestion with HincII produces X-structures corresponding to cross-linked plasmids and linear species corresponding to unhooked plasmids. Right, pICL^Pt^ was replicated in egg extract supplemented with okadaic acid, as indicated. Recovered replication intermediates were digested with HincII and resolved on a denaturing (alkaline) agarose gel. Intermediates were detected by Southern blotting with ^32^P radiolabeled probes and visualized by autoradiography. Samples were run alongside unreplicated pICL^Pt^ that was digested with HincII and BamHI or AflIII to generate standards corresponding to cleavage of both parental DNA strands on one side or the other of the ICL (referred to as long or short *cis*-double linear). Note that replication in the presence of okadaic acid results in an accumulation of species that comigrate with both types of *cis*-double linear species, indicating that these species are derived from replication intermediates in which both parental DNA strands in one fork have been cleaved. **c**, Purified recombinant hPP2A-AC complex used in Fig. 2c was resolved by SDS-PAGE and visualized with Coomassie stain. **d**, Phosphatase activity of the hPP2A-AC complex shown in **c** was confirmed using the Serine/Threonine Phosphatase Assay System (Promega). Activity was normalized to the maximum for each experiment. Data represent the mean ± standard deviation from three independent experiments. Note that hPP2A-AC phosphatase activity is suppressed to background levels in the presence of okadaic acid or cytostatin.

**Supplementary Figure 3.**
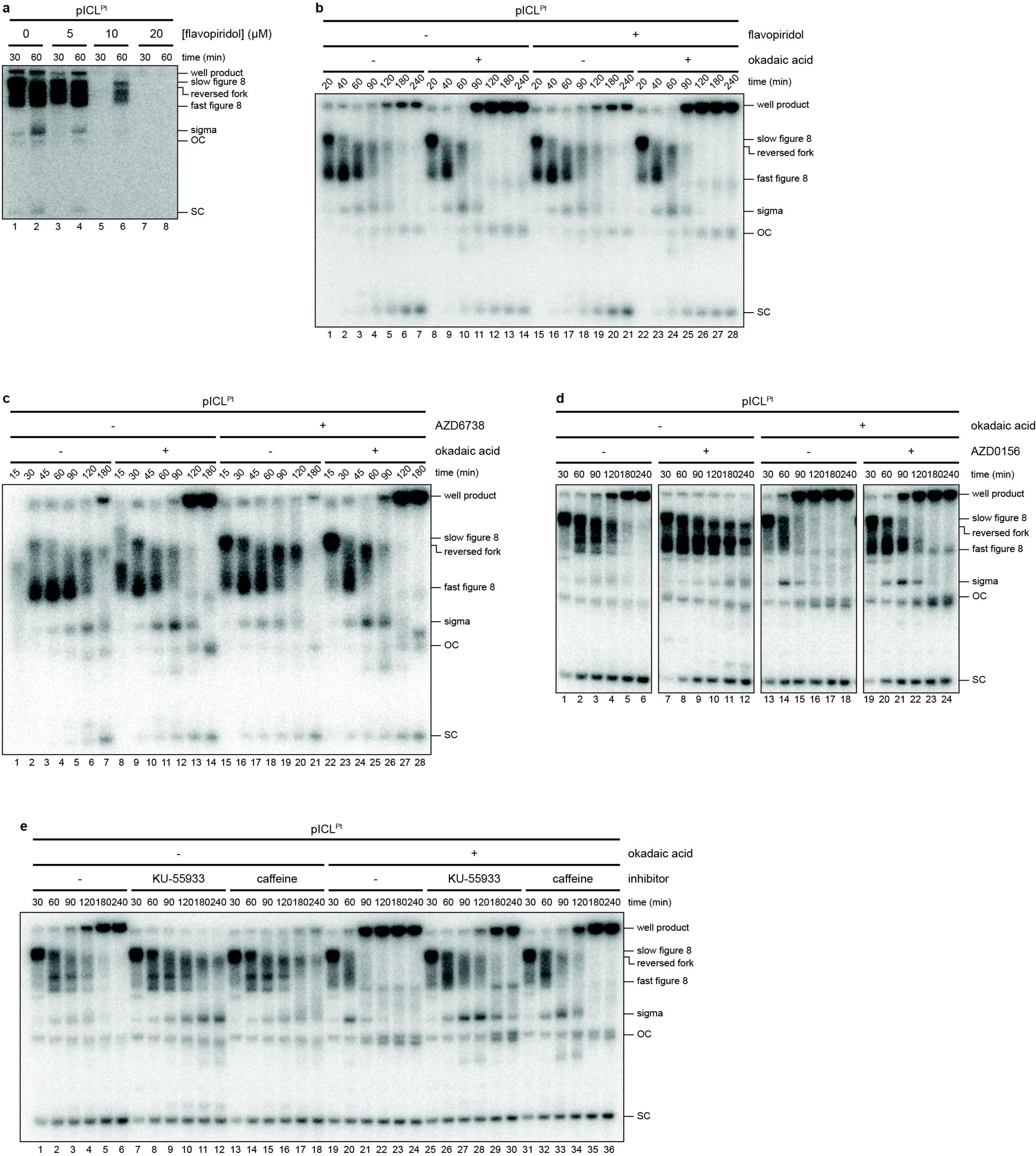
Effects of CDK and DNA damage checkpoint inhibitors on ICL repair by the FA pathway. **a**, pICL^Pt^ was replicated in egg extract supplemented with [α-^32^P]dCTP and pretreated with the pan-CDK inhibitor flavopiridol, as indicated. Replication intermediates were resolved on a native agarose gel as in Fig. 2a. Note that treatment with flavopiridol prior to initiation of replication is sufficient to abolish replication, indicating that the concentration of flavopiridol used in subsequent reactions effectively inhibits CDK activity in egg extracts. **b,** pICL^Pt^ was replicated in egg extract supplemented with [α-^32^P]dCTP and treated with flavopiridol after initiating replication, as indicated. Replication intermediates were resolved on a native agarose gel as in Fig. 2a. Flavopiridol treatment has no discernible effect on processing of ICL repair intermediates, either in the presence or absence of okadaic acid. **c**, pICL^Pt^ was replicated in egg extract supplemented with [α-^32^P]dCTP and the ATR inhibitor AZD6738, as indicated. Replication intermediates were resolved on a native agarose gel as in Fig. 2a. Treatment with AZD6738 results in modest stabilization of reversed fork intermediates but does not suppress okadaic acid-induced formation of putative fork cleavage products or well products. **d**, pICL^Pt^ was replicated in egg extract supplemented with [α-^32^P]dCTP and the ATM inhibitor AZD0156, as indicated. Replication intermediates were resolved on a native agarose gel as in Fig. 2a. As with KU-55933 treatment, AZD0156 treatment resulted in stabilization of reversed fork intermediates and suppressed formation of okadaic acid-induced putative fork cleavage products. **e**, pICL^Pt^ was replicated in egg extract supplemented with [α-^32^P]dCTP and okadaic acid and/or the ATM inhibitors KU-55933 or caffeine, as indicated. Replication intermediates were analyzed as in Fig. 2a.

**Supplementary Figure 4.**
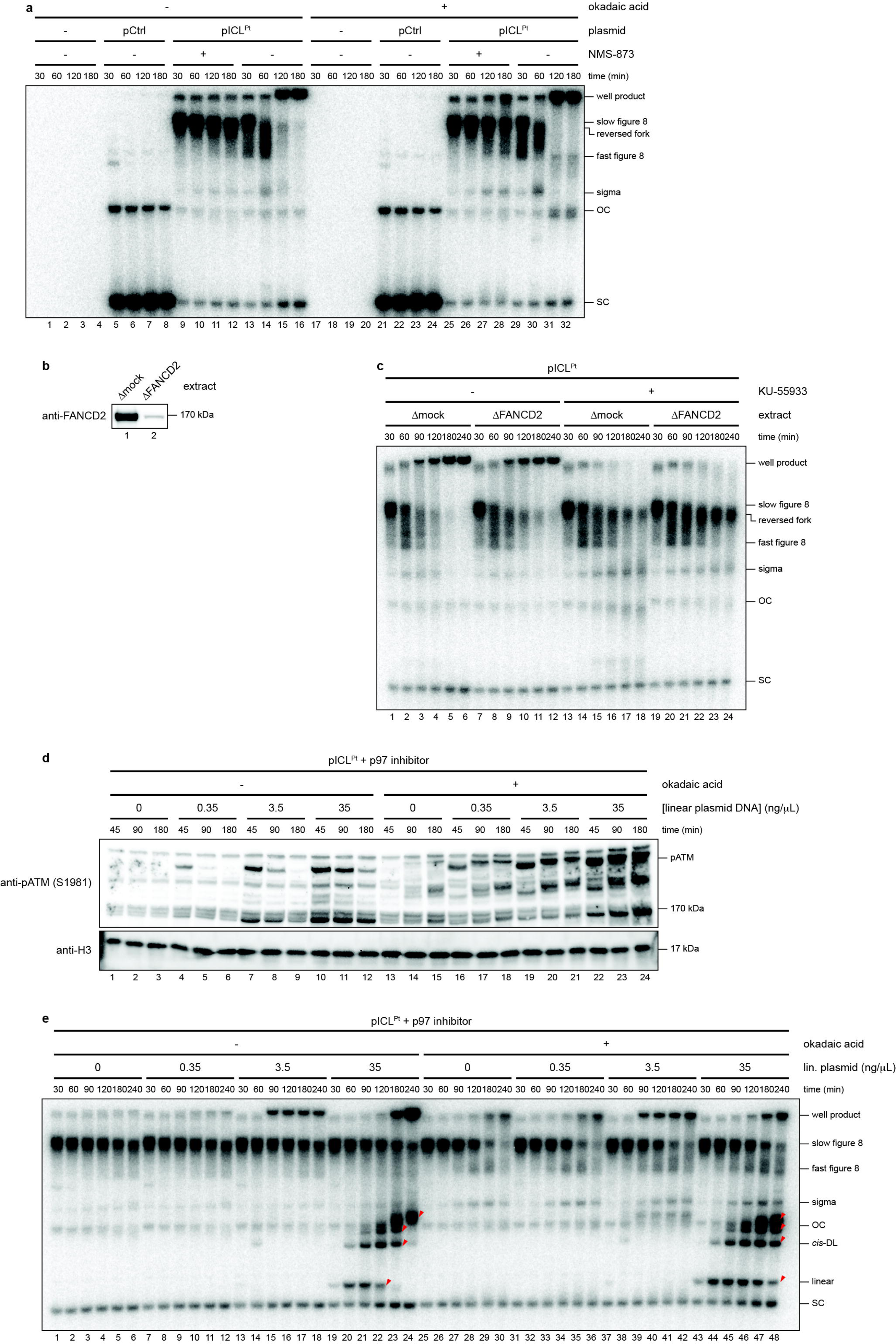
ATM activation promotes processing of reversed fork intermediates. **a**, The replication reactions described in Fig. 4a were supplemented with [α-^32^P]dCTP, and replication intermediates were resolved on a native agarose gel as in Fig. 2a. **b**, FANCD2 immunodepletion. Egg extracts used in Fig. 4b were resolved by SDS-PAGE and blotted for FANCD2. **c**, The replication reactions described in Fig. 4b were supplemented with [α-^32^P]dCTP, and replication intermediates were resolved on a native agarose gel as in Fig. 2a. **d**, pICL^Pt^ was replicated in egg extract supplemented with NMS-873 (to prevent CMG unloading), okadaic acid, and increasing concentrations of linearized pCtrl, as indicated. Replication reactions were blotted for phospho-ATM (S1981) and H3 as in Fig. 3b. Addition of linear DNA induces robust ATM activation. **e**, The pICL^Pt^ replication reactions described in **d** were additionally supplemented with [α-^32^P]dCTP, and replication intermediates were resolved on a native agarose gel as in Fig. 2a. Despite extensive ATM activation, slow figure 8 converged replication fork structures are largely refractory to resection and fork collapse. Red arrowsheads indicate linear plasmids and end-joining products that become labeled due to unscheduled DNA synthesis in extract.

**Supplementary Figure 5.**
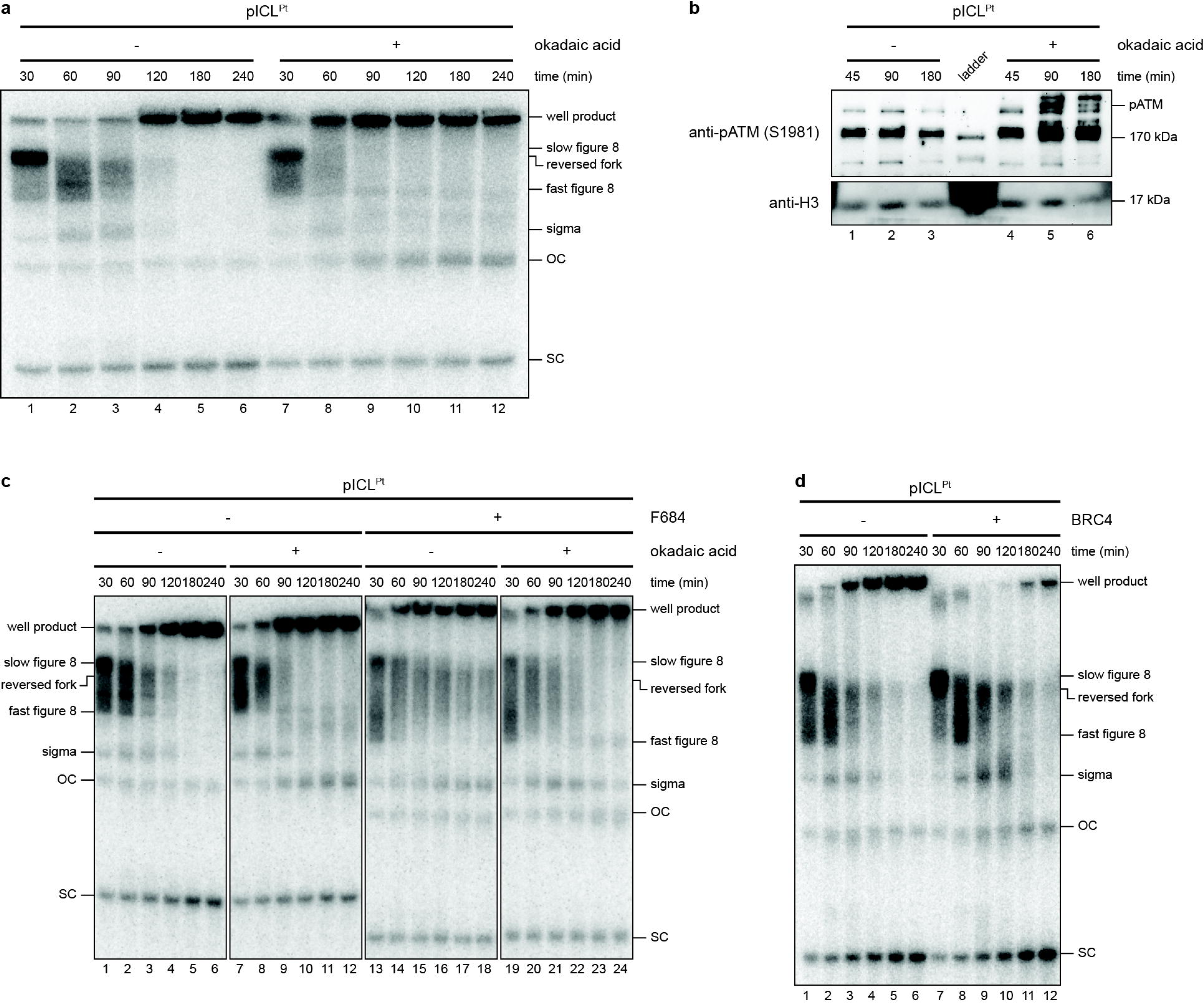
Nuclease-dependent processing of reversed fork intermediates. **a**, The untreated and okadaic acid-treated replications analyzed by mass spectrometry in Fig. 5a were supplemented with [α-^32^P]dCTP, and replication intermediates were resolved on a native agarose gel as in Fig. 2a. **b**, The untreated and okadaic acid-treated replications analyzed by mass spectrometry in Fig. 5a were resolved by SDS-PAGE and blotted for phospho-ATM (S1981) and H3 to confirm ATM activation. **c**, pICL^Pt^ was replicated in egg extract supplemented with [α-^32^P]dCTP and okadaic acid and the EXO1 inhibitor F684, as indicated. Replication intermediates were analyzed as in Fig. 2a. Note that the replication reactions shown were performed in parallel, but lanes 1-12 and lanes 13-24 were resolved on separate native agarose gels, resulting in differences in the migration of replication intermediates and products across gels. **d**, pICL^Pt^ was replicated in egg extract supplemented with [α-^32^P]dCTP and BRC peptide, as indicated. Replication intermediates were analyzed as in Fig. 2a. BRC peptide prevents RAD51 filament formation. Addition of BRC peptide suppressed the formation of well products, consistent with abrogation of RAD51 filament formation and inhibition of homologous recombination. However, BRC peptide treatment did not destabilize reversed fork intermediates or trigger formation of fork cleavage products, indicating that compromised RAD51-dependent fork protection is not sufficient to induce over-processing of reversed fork intermediates.

## Acknowledgements

We thank J. Campbell and members of the Semlow lab for comments on the manuscript. We thank I. Fianu for providing the PP2A expression construct and B. Shen for providing EXO1 inhibitors. D.R.S. is supported by NIH grant no. R01GM151410 and a Shurl and Kay Curci Foundation Research Grant and is a Ronald and JoAnne Willens Scholar and a Vallee Scholar.

## Author contributions

M.V.A. performed most of the experiments using egg extracts. V.A.M. performed the CRISPR dropout screens described in Fig. 1b,c and Supplementary Fig. 1 as well as the experiments described in Fig. 2c, Fig. 5e, and Supplementary Fig. 1a–c. J.T. performed the experiments described in Fig. 1d,e. R.N. purified and determined the activity of the recombinant hPP2A-AC complex described in Supplementary Fig. 2c,d. T.Y.W., and T.F.C. assisted with performing and analyzing the experiment described in Fig. 5a. M.V.A., V.A.M. and D.R.S. designed the experiments. M.V.A. and D.R.S. wrote the manuscript with input from all authors.

## Methods

### Cell lines

RPE1-hTERT Cas9 *TP53^-/-^* were a gift from the Durocher lab and generated as described^65^. RPE1-hTERT Cas9 *TP53^-/-^ FANCD2^-/-^* were generated by nucleofecting RPE1-hTERT Cas9 *TP53^-/-^* cells with a plasmid containing the sgRNA sequence targeting FANCD2 (5’-ATTCCCAGCACGCTGATGTG-3) using a custom nucleofection buffer (25 mM Na_2_HPO_4_ [pH 7.75], 2.5 mM KCl, 11 mM MgCl_2_) and Lonza nucleofection cuvettes in an Amaxa 4D nucleofector using program CN114. 24 hours after transfection, cells were selected with gentamicin (Thermo Fisher Scientific #15-710-064). After an additional 72 hours, single cells were sorted into 96-well plates on a BD FACSARIA II Cell sorter instrument and grown until colonies formed. *FANCD2^-/-^* clones were selected based on successful gene editing determined by PCR amplification and TIDE analysis^66^, and the knockout phenotype was confirmed via immunoblotting for FANCD2. RPE1-hTERT-derived cell lines were grown in Dulbecco’s Modified Eagle medium (DMEM; Gibco #11965126) supplemented with 10% heat inactivated fetal bovine serum (Gibco #10082147) and 2 µg/mL blasticidin (Invivogen #ant-bl-05). HEK Lenti-X 293T (Takara #632180) cells were grown in DMEM (Gibco #10569010). All cell lines were grown at 37 °C, 5% CO_2_. All cell lines routinely tested negative for mycoplasma using a Universal Mycoplasma Detection Kit (American Type Culture Collection #30-1012K) or a MycoAlert Mycoplasma Detection Kit (Lonza #LT07-418) with the MycoAlert Assay Control Set (Lonza #LT07-518) per manufacturers’ protocols.

RPE1-hTERT Cas9 *TP53^-/-^ PTPA^-/-^* and RPE1-hTERT Cas9 *TP53^-/-^ FANCD2^-/-^ PTPA^-/-^* cell lines were generated using the CRISPR-Cas9 system. Oligonucleotides encoding guide RNAs targeting PTPA (5’-ACAGCAGAGGAAAGCAGCG) and a negative control (5’-GGGGCCACTAGGGACAGGAT) were cloned into the lentiCRISPRv2 vector, (Addgene #52961). Lentivirus was produced by seeding HEK293T cells at 60% confluence in 6-well plates 24 hours before transfection. The gRNA-lentiCRISPRv2 construct was transfected together with the packaging plasmids (pol/rev, VSVg, tat) using X-tremeGENE 9 DNA transfection reagent (Millipore Sigma #6365779001). Virus was harvested 48 hours after transfection and filtered through a 0.45 µm membrane. Virus was transduced into *FANCD2^+/+^* or *FANCD2^-/-^* cells with 8 µg/mL polybrene (Millipore Sigma #tr-1003-g) at a density of 10%. After 48 hours, puromycin selection was performed with 25 µg/mL puromycin (Invivogen #ant-pr-1) for 72 hours. Knockouts were validated by immunoblotting.

### CRISPR screens

CRISPR screens were performed as previously described^67^, with some modifications. RPE1-hTERT Cas9 *TP53^-/-^* and RPE1-hTERT Cas9 *TP53^-/-^ FANCD2^-/-^* cells were transduced with the lentiviral TKOv3 library (Addgene #125517) at a low MOI (0.3), and puromycin-conditioned media was added 24 hours following transduction. The following day, cells were trypsinized and replated in the same plates while maintaining the puromycin selection. The puromycin selection was maintained for an additional 48 hours following replating. Four days after transduction, which was considered the initial time point (t0), cells were pooled and divided into two technical replicates. Screens were performed by subculturing cells at days 3 and 6 (t3 and t6), at which point each replicate was divided into untreated and treated conditions. Cells were subcultured in medium without drug every three days (t9, t12, and t15). UVA and psoralen/UVA (PUVA) treatments were applied one day after subculturing cells (t7, t10, t13 and t16). PUVA treatment was applied according to previous determination of LD20 concentrations in uninfected cells. For PUVA treatment, plates were washed with PBS and the media replaced with Hanks’ Balanced Salt Solution (HBSS), calcium, magnesium (Gibco #24020117) either containing DMSO or trimethylpsoralen (Millipore Sigma #6137) and incubated in the dark, under normal atmospheric conditions for 45 minutes. After incubation, plates were irradiated with UVA 4 × 200,000 µJ, rotating the plate 45 degrees between every dose for even coverage. After incubation, conditioned HBSS was aspirated and replaced with fresh DMEM. On t18, cell pellets were frozen for genomic DNA extraction. Screens were performed in technical duplicates and library coverage of >400 sgRNA was maintained at every passage. gDNA from cell pellets was extracted using the QIAamp Blood Maxi Kit (QIAGEN # 51194), and genome-integrated sgRNA sequences were amplified via PCR using NEBNext Ultra II Q5 Master Mix (New England Biolabs [NEB] #M5044) using primers forward: 5’-GAGGGCCTАТТССCATGATTC and reverse: 5’-CAAACCCAGGGCTGCCTTGGAA^67^. A second PCR reaction containing i5 and i7 multiplexing barcodes was completed, and products were gel purified and evaluated with an Agilent BioAnalyzer using the High Sensitivity Kit (Agilent #5067-4626) before sequencing on Illumina NextSeq2000 to determine sgRNA representation in each sample.

### Quantification, statistical analysis, and data quality control

The sgRNA read count files were computed from the raw CRISPR fastq files using the ‘count function from the MAGeCK software^68^. The gene scores (normZ values) were estimated from the read count files using the DrugZ algorithm, testing the PUVA treated replicates against their respective non-treated controls. The MAGeCK count summary provides a summary to assess screen data quality. Both screens had a mapped reads percentage above 86% and Gini indexes below 0.095.

### Cell viability assay

FANCD2^+/+^ PTPA^+/+^, FANCD2^+/+^ PTPA^-/-^, FANCD2^-/-^ PTPA^+/+^, and FANCD2^-/-^ PTPA^-/-^ cells were plated at 2000 cells per well in a 96-well plate in triplicate. 2 mM cisplatin stock solution was made by dissolving cis-Diammineplatinum(ii) dichloride (Sigma #P4394) in 154 mM NaCl, aliquoted, and and frozen at −80 °C. The stock solution was diluted and titrated down to treat the cells at final cisplatin concentrations of 0, 0.04, 0.08, 0.16, 0.32, 0.63. 1.25, 2.5, 5, and 10 µM. Cells were cultured in treatment for 5 days, after which cell viablity was measured using CellTiter-Glo 2.0 (Promega #G9241).

### Protein expression and purification

Recombinant human PP2A-AC was expressed and purified essentially as previously described^69^. The Bac-to-Bac Baculovirus Expression System (Thermo Fisher Scientific #10359016) was used to generate baculoviruses expressing 6xHis-PP2A-AC according to the manufacturer’s protocols. Hi5 insect cells were transfected with the baculoviruses, and recombinant protein was expressed in 2 L of Hi5 suspension culture. Cells were collected once the cell viability dropped to about 80%, generally after 48–72 hours. Harvested cells were lysed by sonication in lysis buffer (50 mM Tris-HCl [pH 8.0], 100 mM NaCl, 20 mM imidazole, 284 µg/mL leupeptin, 1.37 µg/ml pepstatin A, 0.17 mg/mL phenylmethylsulfonyl fluoride [PMSF], 0.33 mg/mL benzamidine, 1X Roche cOmplete EDTA-free Protease Inhibitor Cocktail). The lysate was spun at 68,900 x g in an A27-8×50 fixed angle rotor using a Sorvall Lynx 4000 centrifuge, and the soluble fraction was incubated with Ni-NTA resin (Qiagen #30230) overnight at 4 °C. After binding, the resin was washed extensively with wash buffer (50 mM Tris-HCl [pH 8.0], 500 mM NaCl, 50 mM imidazole, 10 mM ATP, 20 mM MgCl_2_, 284 µg/mL leupeptin, 1.37 µg/mL pepstatin A, 0.17 mg/mL PMSF, 0.33 mg/mL benzamidine). His-tagged hPP2A-AC was eluted with elution buffer (50 mM Tris-HCl [pH 8.0], 300 mM NaCl, 200 mM imidazole, 5 mM β-mercaptoethanol [BME], 284 μg/mL leupeptin, 1.37 μg/mL pepstatin A, 0.17 mg/mL PMSF, 0.33 mg/mL benzamidine). The protein was buffer exchanged into storage buffer (50 mM Tris-HCl [pH 8.0], 300 mM NaCl, 1 mM DTT, 10% glycerol), aliquoted, snap frozen, and stored at −80 °C.

### Phosphatase activity assay

Serine/threonine phosphatase activity of recombinant 6xHis-hPP2A-AC was determined using a nonradioactive molybdate dye-based phosphatase assay kit (Promega #V2460) according to the manufacturer’s instructions. 5 µM of protein in assay buffer (250 mM imidazole [pH 7.2], 1 mM ethylenediaminetetraacetic acid [EDTA], 0.1% BME and 0.5 mg/mL bovine serum albumin [BSA]) was incubated with 10 µM okadaic acid or 100 µM cytostatin for 15 mins at room temperature followed by addition of 100 µM substrate phosphothreonine peptide (RRA[pT]VA). The 50 µL reaction was set up in a 96-well plate and incubated at 37 °C for 30 mins. The reaction was stopped by adding 50 µL molybdate dye additive mix, and color was developed by incubating at room temperature for 30 mins. A standard curve was obtained using KH_2_PO_4_ at concentrations ranging from 2 to 40 µM. Absorbance at 600 nm was measured, and the amount of free phosphate released was quantified.

### Preparation of *Xenopus* egg extracts

Animal work was performed at Caltech, which has an approved Animal Welfare Assurance (no. D16-00266) from the NIH Office of Laboratory Animal Welfare. All procedures involving *Xenopus laevis* were approved by IACUC (Protocol IA20-1797, approved May 28, 2020). Preparation of high-speed supernatant (HSS) and nucleoplasmic extract (NPE) from *Xenopus laevis* oocytes was done essentially as described previously^70^. In brief, six laboratory-bred wild-type adult female *X. laevis* (aged >2 years) were primed with 40 IU of human chorionic gonadotropin (hCG), followed by an injection of 500 IU hCG 2–8 days later. Eggs were collected the next day, dejellied in 1 L 2.2% (w/v) cysteine, pH 7.7, and washed first with 2 L 0.5X Marc’s Modified Ringer’s (MMR) solution (2.5 mM HEPES-KOH [pH 7.8], 50 mM NaCl, 1 mM KCl, 0.25 mM MgSO_4_, 1.25 mM CaCl_2_, 0.05 mM EDTA) and then with 1 L Egg Lysis Buffer (ELB) (10 mM HEPES-KOH [pH 7.7], 50 mM KCl, 2.5 mM MgCl_2_, 250 mM sucrose, 1 mM dithiothreitol [DTT], and 50 μg/mL cycloheximide). Eggs were transferred to 14 mL round bottom Falcon tubes (Corning #352059) and packed by spinning at 200 x g for 1 minute using a Sorvall ST 8 centrifuge (Thermo Fisher Scientific). Apoptotic eggs were aspirated from each tube, and the eggs were supplemented with 5 µg/mL aprotinin, 5 µg/mL leupeptin, and 2.5 µg/mL cytochalasin B before being crushed by centrifugation at 20,000 x g for 20 mins at 4 °C in a TH13-6×50 swinging bucket rotor using a Sorvall Lynx 4000 centrifuge. Low-speed supernatant (LSS) was collected, supplemented with 50 μg/mL cycloheximide, 1 mM DTT, 10 μg/mL aprotinin, 10 μg/mL leupeptin, and 5 μg/mL cytochalasin B, and spun at 260,000 x g for 90 mins at 2 °C in thinwall ultra-clear centrifuge tubes (Beckman Coulter #347356) in a TLS-55 swinging bucket rotor using an Optima MAX-E tabletop ultracentrifuge (Beckman Coulter). The lipids that accumulated at the top of each tube were aspirated, and the cytosolic layer underneath was recovered, aliquoted, snap frozen in liquid nitrogen, and stored at −80 °C.

NPE preparation was begun the same way, except that twenty frogs were used rather than six, and the volumes of 2.2% (w/v) cysteine, MMR, and ELB used to dejelly and wash the eggs were doubled. The protocols diverged after LSS collection – for an NPE preparation, the LSS was supplemented with 50 μg/mL cycloheximide, 1 mM DTT, 10 μg/mL aprotinin, 10 μg/mL leupeptin, 5 μg/mL cytochalasin B, and 3.3 µg/mL nocodazole. The LSS was then transferred to 14 mL Falcon tubes (Corning #352059) and spun at 20,000 x g for 15 mins at 4 °C in a TH13-6×50 swinging bucket rotor using a Sorvall Lynx 4000 centrifuge to remove precipitates. The supernatant was transferred to a 50 mL conical tube and allowed to warm to room temperature, at which point it was supplemented with 2 mM ATP, 20 mM phosphocreatine, and 5 μg/mL creatine phosphokinase. Demembranated sperm chromatin was gradually added to a final concentration of 6,600 U/µL to initiate nuclei formation, which was allowed to proceed for 75–90 mins before nuclei were separated by spinning the extract at 18,000 x g for 3 mins at 4 °C in glass culture tubes (VWR #47729-572) in a TH13-6×50 swinging bucket rotor using a Sorvall Lynx 4000 centrifuge. The nuclei that accumulated at the top of each tube were harvested, transferred to either thickwall (Beckman Coulter #347778) or thinwall (Beckman Coulter #347356) ultracentrifuge tubes (depending on the volume of nuclei collected), and spun at 260,000 x g for 30 mins at 2 °C in a TLS-55 swinging bucket rotor using an Optima MAX-E tabletop ultracentrifuge (Beckman Coulter). The lipids that accumulated at the top of each tube were aspirated, and the nucleoplasmic layer underneath was recovered, aliquoted, snap frozen in liquid nitrogen, and stored at −80 °C.

### Immunodepletions

Immunodepletions using antibodies raised against FANCD2 were performed essentially as described previously^31^. Briefly, Protein A Sepharose Fast Flow (Cytiva #17127903) resin was washed four times with 1X phosphate-buffered saline (PBS) by spinning at 622 x g for 30 seconds in an S-24-11-AT rotor using an Eppendorf 5430R centrifuge. These spin conditions throughout this depletion protocol, unless stated otherwise. After the washes, the resin incubated with 3.6 volumes of anti-FANCD2 serum overnight at 4 °C on a rotating wheel. The next day, the resin was washed twice with 1X PBS, once with ELB mix (2.5 mM MgCl_2_, 50 mM KCl, 10 mM HEPES, 250 mM sucrose), twice with ELB mix supplemented with 0.5 M NaCl, and three times with ELB mix. Three rounds of depletion were performed by adding one volume of HSS or NPE to 0.14 volumes of resin and incubating on a rotating wheel for 20 mins per round at room temperature. After the final round, extract supernatants were collected and used for replication reactions, as described below.

### Preparation of oligonucleotide duplexes with site-specific interstrand cross-links

Cross-linked oligonucleotides were prepared essentially as previously described^4,6,71^. To generate an oligonucleotide duplex containing a cisplatin-ICL, 1 mM cisplatin was incubated in 10 mM NaClO_4_ (pH 5.2), 0.95 mM AgNO_3_ for 24 hours at 37 °C in the dark to produce monoaquamonochloro cisplatin. Then, the cisplatin monoadduct was produced by incubating 0.125 mM Pt-ICL top oligonucleotide with 0.375 mM monoaquamonochloro cisplatin in 5.63 mM NaClO_4_ (pH 5.2) for 12 mins at 37 °C, after which NaCl was added to 0.1 M to quench the reaction. The top oligonucleotide containing a cisplatin monoadduct was purified from a 20% acrylamide/bis (19:1), 1X TBE, 8 M urea gel by excising the gel slice containing the monoadducted oligonucleotide, crushing it, and eluting into Tris-EDTA (TE) buffer. 1.05 molar equivalents of Pt-ICL bottom oligonucleotide were added to the purified monoadducted top oligonucleotides in 100 mM NaClO_4_, and cross-linking was promoted by incubation at 37 °C for 48 hours. The cross-linked duplex was gel-purified in the same way as the monoadduct. The purified cross-linked oligonucleotide duplex was eluted into 10 mM Tris-HCl (pH 7.4), 10 mM NaClO_4_, snap frozen in liquid nitrogen, and stored at −80°C.

To generate the oligonucleotide duplex containing an ICL formed through an abasic (AP) site, the corresponding top and bottom oligonucleotides (AP-ICL top and AP-ICL bottom) were annealed at a concentration of 5 µM each in 30 mM HEPES-KOH (pH 7.0), 100 mM NaCl by heating to 95 °C for 5 mins and cooling at 1 °C/min to 18 °C on a Bio-Rad T100 Thermal Cycler. The annealed duplex was then digested with 0.05 U/µL uracil-DNA glycosylase (NEB #M0280) in 1X UDG buffer (20 mM Tris-HCl, 10 mM DTT, 10 mM EDTA [pH 8.0]) for 120 mins at 37 °C, extracted with phenol:chloroform:isoamyl alcohol (25:24:1; pH 8.0), precipitated in ethanol, and resuspended in 50 mM HEPES-KOH (pH 7.0), 100 mM NaCl. The cross-link was formed by incubating the oligonucleotide duplex at 37 °C for 5–7 days, after which the DNA was precipitated in ethanol, suspended in formamide loading dye (86% formamide, 1X TBE, 20 mM EDTA, 0.025% [w/v] xylene cyanol, and 0.025% [w/v] bromophenol blue), and purified from a 20% acrylamide/bis (19:1), 1X TBE, 8 M urea gel as described above. The cross-linked duplex was eluted into TE buffer, extracted with phenol:chloroform:isoamyl alcohol (25:24:1; pH 8.0), precipitated in ethanol, resuspended in 10 mM Tris-HCl (pH 8.5), aliquoted, snap frozen in liquid nitrogen, and stored at −80 °C.

### Preparation of interstrand cross-link containing plasmids (pICL)

Plasmids containing site-specific ICLs were prepared exactly as described previously^72^.

### Replication reactions

Plasmids were replicated in *Xenopus* egg extracts at a final concentration of 2.5 ng/µL (except where indicated otherwise) essentially as described previously^73^. Where indicated, licensing mix was supplemented with 167-333 nM 3000 Ci/mmol [α-^32^P]dCTP (Revvity #BLU513H250UC). Where indicated, NPE mix was supplemented with 300 µM NMS-873 (Millipore Sigma #SML1128), 7.5 mM caffeine (Millipore Sigma # C0750), 7.5–30 µM flavopiridol (MedChemExpress #HY-10005), 4 µM AZD6738 (MedChemExpress #HY-19323), or 75 µM BRC4 (Biosynth #CRB1001302). Replication was initiated by mixing 1 volume of licensing mix with 2 volumes of NPE mix. Where indicated, replication reactions were supplemented with 1–6 µM okadaic acid (Tocris Bioscience #1136) or 250 µM cytostatin (Santa Cruz Biotechnology #sc-394496) 5 minutes into the reaction; 300 µM KU-55933 (MedChemExpress #HY-12016) or 100 µM AZD0156 (MedChemExpress #HY-100016) 20 minutes into the reaction; or 0.35–35 ng/µL linear plasmid, 4 µM hPP2A-AC, 5 mM C5 (MedChemExpress #HY-128729), 3 mM C200, 300 µM F684, 20 µM flavopiridol, or 1 mM HRO761 (Selleckchem #E1908) 30 minutes into the reaction. For native agarose gel analysis, 1 µL samples of each reaction were quenched into 6 µL replication stop mix (80 mM Tris-HCl [pH 8.0], 8 mM EDTA, 0.13% phosphoric acid, 10% ficoll, 5% sodium dodecyl sulfate [SDS], 0.1% bromophenol blue) at the indicated time points. Samples were worked up and visualized by phosphorimaging as described previously^72^. For Southern blots and nascent strand analysis, 4 µL samples of each reaction were quenched into 40 µL clear stop mix (50 mM Tris-HCl [pH 8.5], 0.5% SDS, 25 mM EDTA) at the indicated time points.

Once all time points were quenched, samples were digested with 200 µg/mL RNAse A for 30 mins at 37 °C and with 1 mg/mL proteinase K for 60 mins at 37 °C. Samples were adjusted to 200 µL with 10 mM Tris-HCl pH 8.5, extracted twice with phenol:chloroform:isoamyl alcohol (25:24:1; pH 8.0), ethanol precipitated, and resuspended in 8–10 µL 10 mM Tris-HCl pH 8.5. Samples were subsequently digested and analyzed as described below for their respective assays.

### Southern blotting

Southern blotting was performed exactly as described previously^72^.

### Nascent strand analysis

Nascent strand analysis was performed essentially as described previously^11^. Briefly, plasmids were replicated and purified as described above. For leading strand analysis, 2 µL of recovered DNA was digested with 2 U AflIII (NEB #R0541) and 2 U of either BamHI (NEB #R0136) or EcoRI-HF (NEB #R3101) at 37 °C for 2.5–4 hours. For lagging strand analysis^74^, 2 µL DNA was digested with 2 U Nt.BstNBI (NEB #R0607) at 37 °C for 2.5–4 hours. Denaturing formamide gel loading buffer (Invitrogen #AM8547) was added 1:1 to each reaction, and nascent strands were resolved on a 7% acrylamide/bis (19:1), 8 M urea, 0.8X GTG buffer (71 mM Tris-HCl, 23 mM taurine, 0.4 mM EDTA) gel. Gels were dried and visualized by phosphorimaging.

### Plasmid pulldown

Plasmid pulldowns were performed essentially as previously described^10^. In brief, 4.5 µL Dynabeads Protein A beads (Invitrogen #10001D) per time point were washed 3 times with wash buffer (50 mM Tris-HCl [pH 7.5], 150 mM NaCl, 1 mM EDTA [pH 8.0], 0.02% Tween 20) on a magnetic rack and bound to 2 pmol biotinylated LacI per µL of beads for 40 mins at room temperature on a rotating wheel. Following the incubation, dynabeads were washed 4 times with pulldown buffer (2.5 mM MgCl_2_, 50 mM KCl, 10 mM HEPES, 250 mM sucrose, 0.25 mg/mL BSA, 0.02% Tween 20) on a magnetic rack, aliquoted for individual time points, and stored on ice. Replication reactions were performed as described above, but at a plasmid concentration of 5 ng/µL in extracts. At the indicated times, 6 µL of each reaction were quenched into 30 µL Dynabeads and incubated on a rotating wheel for 30 mins at 4 °C. Following the incubation, the beads were washed 3 times with 2.5 mM MgCl_2_, 50 mM KCl, 10 mM HEPES, 0.25 mg/mL BSA, 0.03% Tween 20 on a magnetic rack, suspended in 1X Laemmli loading buffer, and analyzed by immunoblotting. 1 µL of each replication reaction was also quenched directly into 1X Laemmli loading buffer to serve as an input sample.

### Immunoblotting

Samples in 1X Laemmli loading buffer, generally corresponding to 0.3-0.75 µL of replication reaction (except for plasmid pulldowns, which are described above), were resolved on 4– 15% (Bio-Rad #5671085, #4561086) or 10% (Bio-Rad #5671035, #4561036) acrylamide gels and transferred to 0.45 µM polyvinyl difluoride (PVDF) membrane (Thermo Fisher Scientific #88518). Membrane was blocked with 5% nonfat milk in 1X PBS with 0.05% Tween 20 (PBST) for 60 mins at room temperature, rinsed with 1X PBST, and incubated in primary antibody in 1X PBST overnight at 4 °C. Commercial primary antibody identities and dilutions used for blotting cell samples were as follows: rabbit polyclonal anti-PTPA (human), Cell Signaling Technology #3330, 1:1,000; mouse monoclonal anti-FANCD2 (human), Santa Cruz Biotechnology #sc-20022, 1:1,000; and mouse monoclonal anti-RPA (human), Millipore Sigma #MABE285, 1:1,000. Commercial primary antibody identities and dilutions used for blotting *Xenopus* egg extract samples were as follows: rabbit monoclonal anti-phospho-ATM (S1981; human), Rockland Immunochemicals #200-301-500, 1:1,500; rabbit polyclonal anti-H3 (human), Cell Signaling Technology #9715, 1:500; rabbit polyclonal anti-phospho-CHK1 (S345; human), Cell Signaling Technology #2341, 1:500; rabbit polyclonal anti-phospho-RPA32 (S33; human), Bethyl Laboratories #A300-246A, 1:1,000; and mouse monoclonal anti-RAD51, Novus Biologicals #NB100-148, 1:1,000.

Rabbit polyclonal anti-CHK2^75^, rabbit polyclonal anti-FANCD2^11^, and rabbit polyclonal anti-MRE11^76^ have been previously described. The next day, membrane was washed 3 times with 1X PBST and incubated in secondary antibody diluted 1:25,000 in 5% nonfat milk in 1X PBST for 30 mins at room temperature. Secondary antibody was either goat polyclonal anti-rabbit IgG (H+L) conjugated to horseradish peroxidase (HRP), Jackson ImmunoResearch #111-035-003 or goat polyclonal anti-mouse IgG (H+L) conjugated to HRP, Jackson ImmunoResearch #115-035-003. Following the secondary antibody incubation, membranes were washed 3 times with 1X PBST, incubated for 2–4 mins in chemiluminescence substrate (either Thermo Fisher Scientific #34076 or Genesee Scientific #20-300S), and imaged using a ChemiDoc Imaging System (Bio-Rad).

### Sample preparation for phosphoproteomics

Plasmids were replicated at a final concentration of 5 ng/µL in *Xenopus* egg extract in the absence or presence of okadaic acid as described above. After 90 minutes of replication, 40 µL samples of each reaction were quenched in triplicate into 400 µL SDS solubilization buffer (5% SDS, 50 mM triethylammonium bicarbonate [TEAB; pH 8.5]) each. Proteins were reduced by adding tris(2-carboxyethyl)phosphine (TCEP) to 5 mM and incubating at 55 °C for 20 mins. After samples cooled to room temperature, proteins were alkylated by adding chloroacetaldehyde (CAA) to 100 mM and incubating in the dark for 30 mins. Samples were spun at 13,000 x g for 8 mins in an Eppendorf 5424R microcentrifuge to remove undissolved matter, and trifluoroacetic acid (TFA) was added to 1%. Sample volume was increased approximately 7-fold by adding S-Trap buffer (90% methanol, 100 mM TEAB [pH 7.55]), and samples were loaded onto S-Trap Midi Columns (ProtiFi). Protein cleanup, on-column digestion with 50 µg trypsin per column, and peptide elution was performed according to the manufacturer’s instructions, and eluted peptides were dried by SpeedVac.

Peptides were reconstituted in 1 mL 1% TFA. Sep-Pak tC18 cartridges (Waters Corp #WAT054960) were placed into 15 mL conical tubes and activated by adding 1 mL 100% acetonitrile (ACN) and spinning at 50 x g for 1 min at room temperature. Note that these spin conditions were also used for all subsequent washes of the Sep-Pak columns. The ACN wash was repeated 3 additional times (4 total washes), after which the columns were equilibrated by washing 4 times with 1 mL 0.1% TFA. The acidified peptide samples were loaded onto the columns and allowed to drain by gravity, and the collected flowthrough was reapplied once. Columns were washed 4 times with 1 mL 0.1% formic acid (FA), and peptides were eluted by gravity 3 times with 350 µL 0.1% FA in 50% ACN. Samples were dried by SpeedVac, after which phosphoenrichment was performed using the Thermo Fisher Scientific TiO_2_ Phosphopeptide Enrichment Kit (#A32993) according to the manufacturer’s instructions. The flowthroughs from the binding and wash steps of the TiO_2_ enrichment were pooled and used for another round of phosphoenrichment with the Thermo Fisher Scientific Fe-NTA Phosphopeptide Enrichment Kit (#A32992) according to the manufacturer’s instructions. Eluates from the two rounds of phosphoenrichment were separately dried by SpeedVac and reconstituted in 0.1% FA.

LC-MS/MS data were acquired in data-independent acquisition (DIA) mode using a Thermo Fisher Scientific Vanquish Neo Tandem UHPLC system coupled to an Orbitrap Astral mass spectrometer. A total of 0.5 µg peptides was injected onto a 75 µm X 2 cm trap column before refocusing on an Aurora UHPLC column (25 cm x 75 µm, 1.7 µm C18, AUR3-25075C18-XT, Ion Opticks) with a flow rate of 0.25 µL/min in a 35-min gradient. The Orbitrap Astral MS was operated at a full MS resolution of 240,000 with a full scan range of 380–980 m/z. The full MS AGC was set to 300%, and IT was set to 10 ms. MS/MS scans were recorded with a 2 Th isolation window, 3.5 ms maximum ion injection time. The isolated ions were fragmented using HCD with 27% NCE.

### Proteomic data analysis

Data search was performed in Proteome Discoverer (PD) (v3.2; Thermo Fisher Scientific) using CHIMERYS^77^ against the 10.1 version of the Xenopus laevis genome downloaded from Xenbase (https://wuehr.scholar.princeton.edu/sites/g/files/toruqf2761/files/xenla_10_1_xenbase.fa sta.txt). Oxidation / +15.995 Da (M) and phosphorylation / +79.9663 (S,T,Y) were set as dynamic modifications, and carbamidomethylation / +57.021 Da (C) was set as a static modification. Downstream analysis was performed using the R packages MSstats (v4.16.1)^78^ and MSstatsPTM (v2.10.3)^63^. Peptide-spectrum match (PSM) tables were imported and processed using the PDtoMSstatsPTMFormat() function. A custom function was implemented to parse Xenbase FASTA headers to extract protein accession identifiers compatible with MSstatsPTM, and peptide abundances were Log_2_-transformed and summarized at the phosphosite and protein levels. Differential abundance analysis between control and okadaic acid-treated samples was performed using linear mixed-effects models, which estimated Log_2_ fold changes (Log_2_FCs) and p-values for each phosphosite. Resulting p-values were adjusted for multiple hypothesis testing using the Benjamini-Hochberg procedure^64^. Phosphosites with FDR-adjusted p-values < 0.05 and absolute Log₂FC > 0.5 were considered significantly regulated.

### Preparation and analysis of next-generation sequencing libraries

Next-generation sequencing (NGS) libraries were prepared and analyzed exactly as described previously^72^. As before, raw sequencing reads were trimmed to 150 bp with Trimmomatic (v0.39)^79^, processed with SIQ (v1.8) using default settings with long-read analysis enabled^80^, and visualized with SIQPlotteR.

## Notes

### Competing Interest Statement

The authors have declared no competing interest.

